# Defining nutrient dependencies of intestinal *Klebsiella pneumoniae* guides rational probiotic engineering

**DOI:** 10.64898/2026.11.02.763509

**Authors:** Ying Zhou, Clement David, Grishma Kulkarni, Tengfei Long, María J. Garzón, Julien Prados, Joachim Kloehn, Olivier Von Rohr, Isabel Meister, Karina B. Xavier, Carles Ubeda, Fangyou Yu, Liang Chen, Simone Becattini

## Abstract

*Klebsiella pneumoniae* is a major cause of antibiotic-resistant infections, yet the ecological determinants enabling its intestinal expansion remain poorly defined. Here, we combined *in vivo* transcriptomics and metabolomics to identify carbon sources exploited by *K. pneumoniae* during colonization of the antibiotic-disrupted gut across strains and dietary contexts. Despite its metabolic breadth, *K. pneumoniae* consistently exploited a restricted set of substrates, including arabinose, raffinose, oligogalacturonate, arabitol, xylose, and gluconate, that contributed to intestinal fitness, with related metabolic signatures also observed in antibiotic-treated human samples. Guided by these findings, we engineered *Escherichia coli* Nissle to enhance consumption of these nutrients and assembled a defined consortium, NICHE, designed to reinforce colonization resistance through nutrient competition. NICHE restricted intestinal expansion of multidrug-resistant and hypervirulent *K. pneumoniae* in preventive and therapeutic settings, with efficacy influenced by dietary nutrient availability. Together, this work defines exploitable *K. pneumoniae* nutrient dependencies and a framework for rational probiotic engineering.

## Introduction

*Klebsiella pneumoniae* (*Kp*) is a major global health threat and a leading cause of antibiotic-resistant infections worldwide. Classified by the WHO as a priority ESKAPE pathogen, *Kp* encompasses both multidrug-resistant (MDR) and hypervirulent (hv) lineages. MDR-*Kp* accounts for a substantial fraction of deaths attributable to antibiotic resistance and is projected to increase sharply in prevalence over the coming decades ^1,2^, prompting the WHO to designate it as the top-priority bacterial pathogen in 2024 ^3^. In parallel, hv-*Kp* causes severe community-acquired infections with high mortality and is increasingly acquiring resistance determinants, generating highly concerning hv-MDR variants ^4^. Together, these trends underscore the urgent need for strategies that prevent *Kp* expansion and transmission rather than relying exclusively on antibiotic treatment.

The gastrointestinal (GI) tract is the primary reservoir for *Kp*. Asymptomatic carriage occurs in a significant fraction of the population ^5–7^ and intestinal abundance is the strongest predictor of subsequent infection by the same strain ^8,9^. Antibiotic exposure impairs colonization resistance provided by the gut microbiota, enabling resident or newly acquired *Kp* strains to expand dramatically and, in some cases, approach near-monocolonization of the intestinal lumen ^5,10–17^. Similar dynamics are observed in mouse models, where *Kp* remains tightly restricted by an intact microbiota but expands rapidly following antibiotic-mediated community disruption ^12,18–22^. These observations indicate that *Kp* expansion in the gut is governed primarily by ecological constraints imposed by competing microbes rather than by host permissiveness.

A central determinant of these constraints is access to carbon. Within the intestine, microbes compete for a diverse but finite pool of dietary- and host-derived substrates. Although the microbiota can restrict *Kp* through mechanisms such as short-chain fatty acid production and immune modulation^23,24^, successful colonization relies largely on *Kp*’s ability to exploit metabolic niches that are inaccessible or underutilized by surrounding microbes. Competition for such carbon sources can restrict pathobiont expansion ^25–27^. Indeed, competition for individual substrates, including xylose ^18^, gluconate ^28^, ethanolamine ^29^ and fucose ^30^ has been shown to limit *Kp* growth in the context of an intact microbiota. In contrast, antibiotic treatment depletes commensal competitors and liberates multiple carbon and nitrogen sources, creating a permissive metabolic landscape that favors *Kp* outgrowth ^20,26^. However, the relative contribution of individual substrates to *Kp* expansion under these conditions remains poorly defined.

Identifying the carbon sources that most strongly support *Kp* colonization is challenging for several reasons. First, *Klebsiella* species possess exceptionally broad metabolic repertoires and can grow on hundreds of distinct nutrients ^31,32^. complicating efforts to identify substrates that are specifically relevant *in vivo*. Second, many carbon utilization pathways are genetically redundant, such that loss of individual genes is buffered by alternative enzymes or parallel routes; consequently, single-gene loss-of-function approaches, including transposon sequencing, often fail to reveal conditionally important metabolic functions ^12,33^. Third, substrate availability in the gut is highly context dependent, varying with diet, microbiota composition, and antibiotic exposure ^17,34^. Finally, the substantial strain-to-strain variation in metabolic preferences raises concerns about the generalizability of colonization determinants identified in individual isolates or conditions ^31,35^.

These limitations have constrained the rational exploitation of metabolic competition as a strategy to control *Kp* colonization. From a translational perspective, metabolic competition offers a promising route to suppress *Kp* without relying on antibiotics. The ability of a competitor to restrict *Kp* expansion depends on metabolic overlap, specifically its capacity to consume the same limiting substrates ^36^. While individual commensals typically provide weak suppression, defined microbial consortia can achieve robust inhibition through coordinated substrate depletion ^26,28,36,37^. However, the clinical deployment of multi-species consortia remains challenging due to safety, manufacturing, and engraftment barriers. An alternative strategy is to engineer a single, well-characterized, aerotolerant organism with broad metabolic interference capacity. *E. coli*, including probiotic strains such as Nissle 1917, has emerged as a partial competitor of *Kp* due to basal metabolic overlap, but its efficacy is inconsistent and context-dependent ^26^.

Here, we used bacterial transcriptomics and fecal metabolomics to define the carbon sources preferentially utilized by *K. pneumoniae* during intestinal colonization in the antibiotic-treated gut across dietary contexts. This approach identified a limited set of substrates that were consistently prioritized despite extensive metabolic redundancy and strain diversity. Notably, these carbon utilization patterns identified in mice were also observed in antibiotic-exposed human samples, suggesting at least partial conservation of *Kp* metabolic dependencies across hosts. Guided by these data, we then constructed a defined collection of *Escherichia coli* strains with complementary substrate utilization profiles designed to deplete these resources *in vivo*, a consortium we named ‘NICHE’ (Nissle Integrated Consortium for Heterogeneous Energy utilization). Together, these results establish carbon source availability as a key determinant of *Kp* intestinal fitness under antibiotic pressure and demonstrate that targeted metabolic competition can be rationally deployed to restrict *Kp* expansion in the gut.

## Results

### *In vivo* transcriptomics identifies carbon utilization pathways associated with *K. pneumoniae* intestinal colonization

We set out to identify the carbon sources utilized by *Kp* during colonization of the antibiotic-depleted intestine, a frequent condition in hospital settings ^10,12^. Given the broad metabolic flexibility of *Klebsiella* species and the presence of strain-specific accessory metabolism that can influence nutrient use and competitive interactions in the gut ^31^, we performed our analyses using two distinct *Kp* strains representing major pathotypes associated with human infection: a multidrug-resistant, carbapenem-resistant strain (MH258) ^12,38^ and a hypervirulent KL2 strain (ATCC 43816) ^39^. Bacteria were grown in LB to either logarithmic or stationary (overnight) phase or introduced into mice that had been treated with antibiotics (ampicillin, metronidazole, neomycin, vancomycin, AMNV) for four days, followed by oral gavage with either strain. Cecal contents were collected one week later. RNA extracted from *in vitro* cultures or cecal samples was subjected to RNA sequencing (**Figure 1A**).

**Figure 1.**
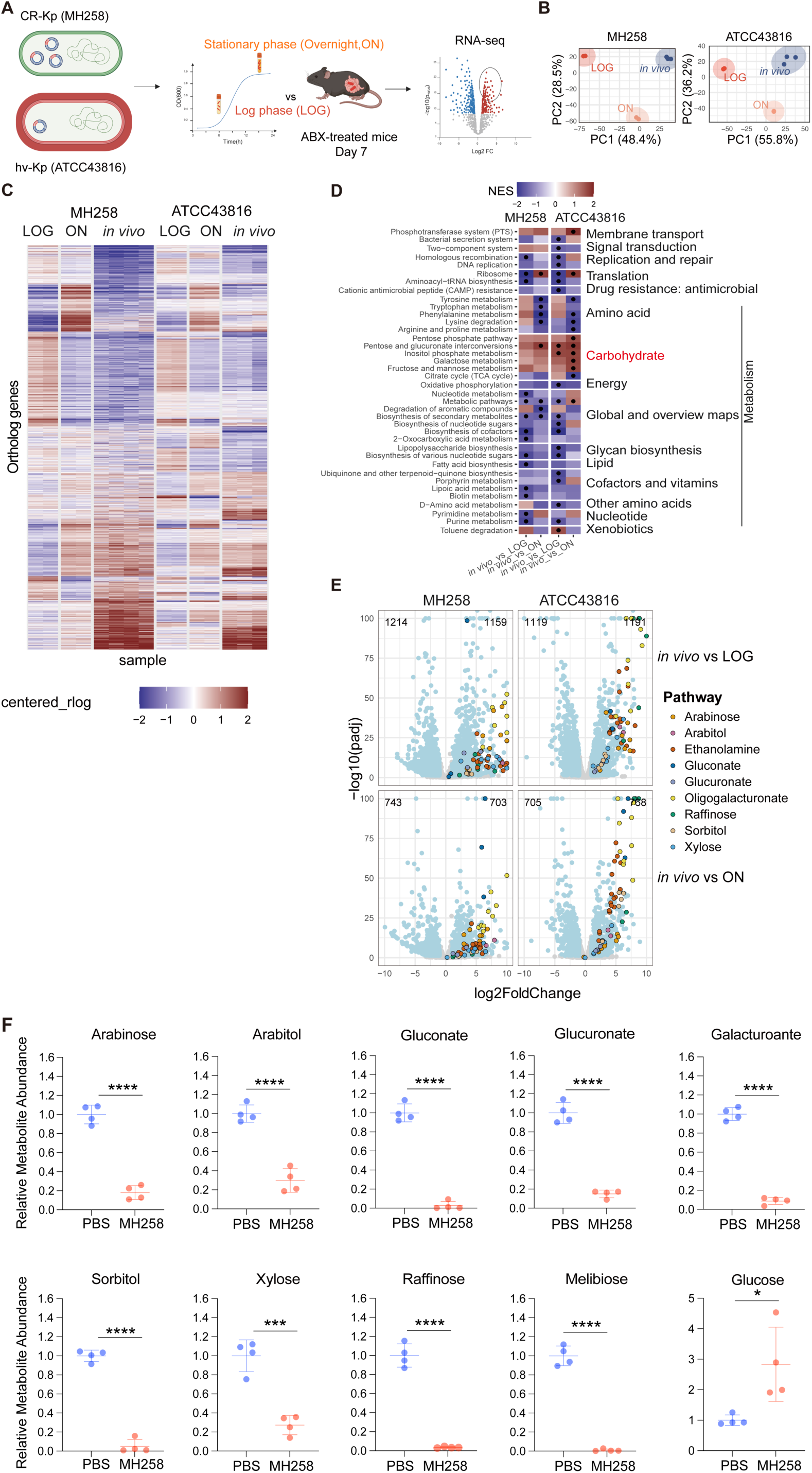
Transcriptome-guided identification of carbon sources supporting *K. pneumoniae* intestinal colonization. **(A)** Schematic overview of the experimental approach. Mice were treated with a broad-spectrum antibiotic cocktail (ampicillin, metronidazole, neomycin, and vancomycin; AMNV)) for 4 days and subsequently colonized by oral gavage with two clinically relevant *K. pneumoniae* strains (MH258, carbapenem-resistant, CR-Kp; ATCC 43816, hypervirulent, hv-Kp). After 7 days, bacterial RNA was isolated from cecal contents or from bacteria grown in LB medium and harvested at logarithmic (LOG) or stationary (overnight, ON) growth phase, and subjected to sequencing. **(B)** Principal component analysis (PCA) of transcriptomes from *in vivo* and *in vitro* samples for both strains. Each dot represents a biological replicate (n=2 for *in vitro* conditions and n=3 for *in vivo* conditions). **(C)** Heatmap and hierarchical clustering of global gene expression profiles from both strains across all conditions. Columns represent individual samples and rows represent genes. Relative gene expression is shown, with red indicating upregulation and blue indicating downregulation compared to *in vitro* conditions. **(D)** KEGG pathway enrichment analysis of genes differentially expressed *in vivo* relative to *in vitro* in both strains. Red indicates pathways enriched among upregulated genes. Black dots denote significantly enriched pathways. **(E)** Volcano plots displaying differential gene expression of *K. pneumoniae in vivo* relative to *in vitro* for both strains. Light blue dots indicate significantly differentially expressed genes (|log2FC| > 1, padj < 0.05), whereas gray dots indicate non-significant genes. Genes associated with metabolic pathways/carbon sources of interest are color-coded. **(F)** GC– MS and LC-MS–based quantification of individual metabolites in the cecal content from ABX (antibiotics)-treated mice one week after reconstitution with *K. pneumoniae*, compared to PBS-treated controls. Each dot represents one mouse (n=4); horizontal lines indicate means ± SD. Note: oligogalacturonate and raffinose cannot be directly detected by GC–MS; melibiose is here used as a proxy for raffinose abundance, as it is generated upon raffinose degradation either enzymatically or during GC–MS processing. Raffinose abundance was quantified directly by LC–MS. Statistical significance was determined using unpaired two-tailed *t* tests (\**P* < 0.05; \*\**P* < 0.01; \*\*\**P* < 0.001; \*\*\*\**P* < 0.0001).

Principal component analysis of the transcriptomes revealed a clear separation between the three growth conditions for both strains (**Figure 1B**). Notably, intestinal *Kp* clustered more closely with stationary-phase than logarithmic-phase bacteria, consistent with reduced nutrient availability and slower growth *in vivo*. When analysis was restricted to genes conserved between the two strains, expression profiles were remarkably similar, indicating shared adaptive strategies across strains despite their genetic divergence (**Figure 1C**). This convergence was also evident at the pathway level: gene set enrichment analysis revealed highly similar transcriptional shifts between *in vitro* and *in vivo* conditions, with the majority of affected pathways related to metabolism. In particular, carbohydrate metabolism was strongly remodeled, indicative of altered carbon source availability in the gut (**Figure 1D**).

To identify metabolic pathways supporting *Kp* intestinal colonization, we focused on genes upregulated *in vivo* relative to each *in vitro* condition, reasoning that induction of metabolic enzymes and transporters would reflect preferred carbon sources in each environment. Strikingly, many of the most strongly upregulated genes were involved in the utilization of specific carbon sources - predominantly sugars - and were conserved across strains and comparisons (**Figures 1E and S1A**). These included genetic modules associated with the metabolism of arabinose, arabitol, ethanolamine, gluconate, glucuronate, oligogalacturonate, raffinose, sorbitol, and xylose, among others. While some of these substrates, such as gluconate ^28^, xylose ^12,18^, and ethanolamine ^29^ have previously been implicated in intestinal colonization by *Klebsiella*, the relevance of the remaining pathways to gut colonization has not been systematically examined (**Figure 1E**). A subset of the identified genes clustered within a conserved chromosomal island present across virtually all *Klebsiella* sequence types, was highly transcribed *in vivo*, and appeared to encode enzymes involved in the utilization of raffinose, oligogalacturonate, and arabinose (**Figure S1B**). These sugars can be derived from pectin, a complex plant cell wall polysaccharide that also contains glucuronate and xylose ^40^ (**Figure S1C**). Consistently, incubation of MH258 with pectin, led to upregulation of genes involved in the metabolism of these sugars, recapitulating the *in vivo* transcriptional profile (**Figures S1D-G**).

Targeted gas chromatography (GC–) and liquid chromatography (LC–) mass spectrometry (MS) confirmed the presence of these carbon sources in the cecal lumen across a range of concentrations (**Figures S2A and B**). Importantly, colonization of antibiotic-treated mice with MH258 or ATCC 43816 led to a marked reduction in the abundance of these sugars, supporting their direct utilization *in vivo* (**Figures 1F and S2C**). In contrast, glucose, which did not emerge as a preferentially utilized substrate from our transcriptomic analysis, was not consumed in *Kp*-colonized mice, suggesting limited *in vivo* utilization despite its canonical role as a preferred bacterial carbon source (**Figures 1F and S2C**), consistent with previous reports ^28^. Together, these data establish transcriptomic profiling as a robust approach to infer *in vivo* carbon source utilization and identify a restricted set of carbon substrates that support *Kp* colonization of the antibiotic-disrupted intestine, including several not previously associated with this process, some of which are likely pectin-derived.

### Transcriptomics-identified metabolic pathways support *in vivo* colonization

To assess the functional relevance of the transcriptomics-identified metabolic pathways for *Kp* intestinal colonization, we generated isogenic deletion mutants in MH258, targeting pathways involved in the utilization of the nine carbon sources highlighted in **Figure 2A**. For arabinose, xylose, glucuronate, and arabitol, we targeted the first enzymatic step of the corresponding catabolic pathways (*araA*, *xylA*, *uxaC*, and *dalD*, respectively). For raffinose and sorbitol, we deleted the corresponding transporters (*rafB,* formerly annotated in this strain as lactose permease *lacY*^12^), and *srlA*; as these deletions encompassed the native promoters, downregulation of downstream pathway genes (in green in the schematic) is expected. For oligogalacturonate utilization, we generated a double knockout strain by deleting both the transporter (*kdgM*) and the first metabolic enzyme (*ogl*), accounting for potential redundancy. For gluconate, we generated two independent mutants, deleting either the transporters (*gntT/U*) or the metabolic enzyme (*gntK*). Finally, to selectively disrupt ethanolamine utilization as a carbon source while preserving ammonia production, we deleted *eutE*, which acts downstream in its catabolic pathway (**Figure 2A**).

**Figure 2.**
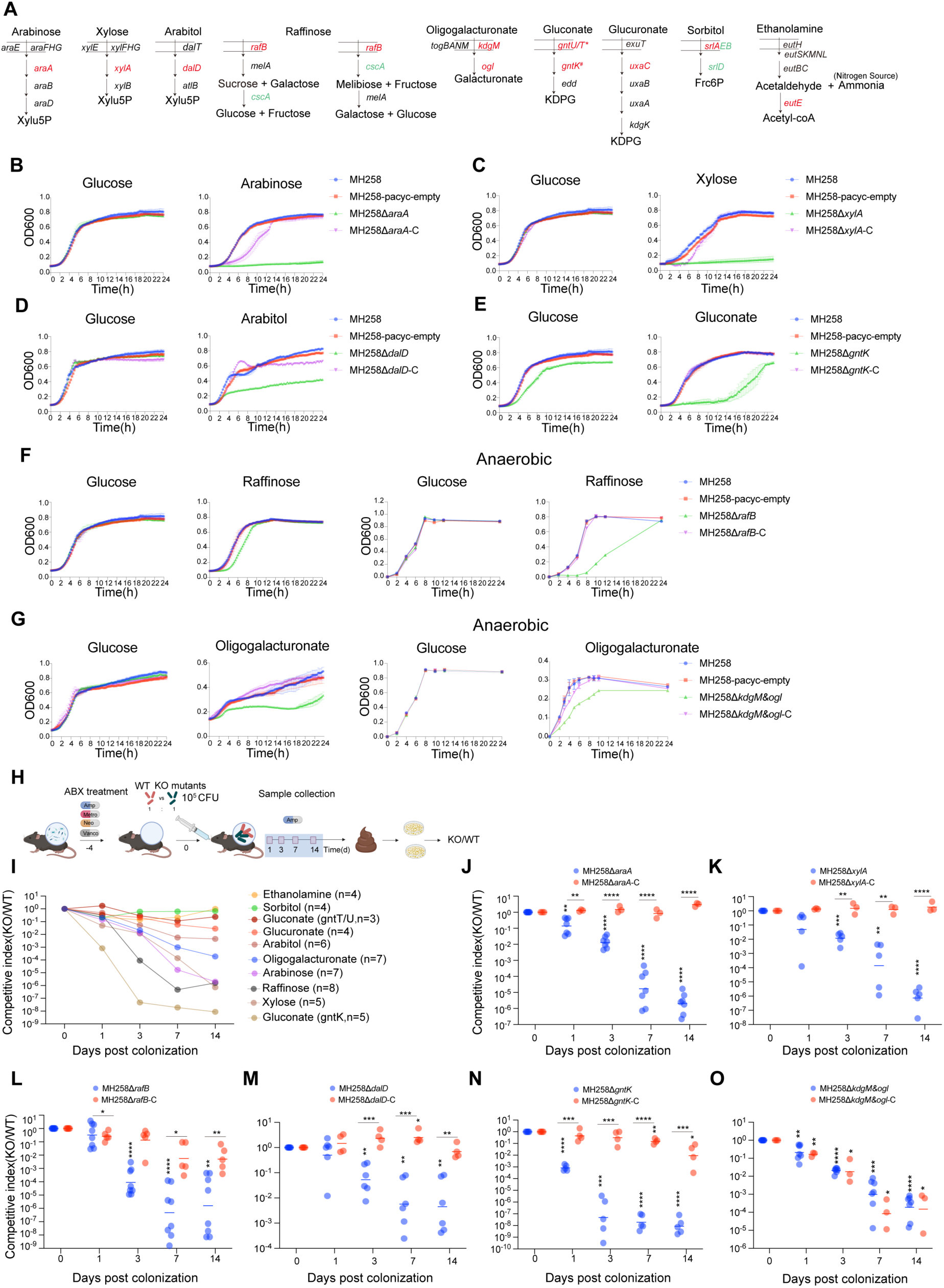
Genetic *in vitro* and *in vivo* validation of candidate carbon utilization pathways contributing to *K. pneumoniae* intestinal colonization. (A) Schematic overview of targeted isogenic deletion mutants constructed, targeting key pathway enzymes (arabinose, xylose, arabitol, glucuronate), combined pathway enzymes/transporters (oligogalacturonate), or operon-level transcriptional knockdown achieved by deletion of the promoter and the first gene of the operon (raffinose, sorbitol). For gluconate utilization, two isogenic deletion mutants were constructed: an asterisk (*) denotes simultaneous deletion of the two gluconate transporters *gntU* and *gntT*, whereas a hash (#) denotes deletion of the key metabolic enzyme *gntK* only. Ethanolamine utilization was selectively blocked at the carbon catabolic level while preserving its role in nitrogen metabolism. Deleted genes are indicated in red and transcriptionally silenced genes in green. (B–G) *In vitro* validation of utilization of specific carbon sources. Growth curves obtained with wild-type, mutant, and complemented strains grown aerobically in minimal M9 medium supplemented with glucose or the selected carbon source. Raffinose-and oligogalacturonate-associated mutants were also assessed under anaerobic conditions. Carbon source concentrations were 0.2% for glucose, xylose, gluconate, and raffinose, and 0.5% for arabitol and oligogalacturonate. Displayed are mean ± SEM values from 3 independent experiments. (H-O) *In vivo* assessment of carbon source contribution to intestinal *K. pneumoniae* colonization. Wild-type and mutant strains were gavaged at a 1:1 ratio into AMNV-treated mice, and fecal samples were collected at indicated time points to calculate competitive indices. Similar experiments were carried out using WT and plasmid-complemented strains. Competitive indices were normalized to the corresponding inoculum ratio and are displayed on a logarithmic scale. A competitive index of 1 indicates no change relative to the inoculum. Each point represents one mouse, n=5-8 mice, from 2–3 independent experiments, and horizontal lines indicate the geometric mean. Statistical analyses were performed on log10-transformed competitive-index values. Symbols above individual groups indicate significance relative to a competitive index of 1, whereas brackets indicate direct comparisons between groups. Comparisons with the neutral value were performed using two-sided one-sample t-tests, and direct comparisons between mutant and complemented mutant groups were performed using two-sided unpaired Welch’s t-tests (\**P* < 0.05; \*\**P* < 0.01; \*\*\**P* < 0.001; \*\*\*\**P* < 0.0001).

We first assessed mutant fitness *in vitro* by measuring growth in minimal media supplemented with the relevant carbon source. For nearly all mutants, growth in glucose was indistinguishable from wild type (WT), indicating intact metabolic fitness, but defective when cultured on the corresponding carbon source (**Figures 2B-G and S3A**). The only exception was the *gntK* mutant, which exhibited delayed growth in glucose and in multiple other sugars, suggesting broader metabolic impairment (**Figures 2E and S3B**). In contrast, a *gntT/U* transporter mutant displayed a more gluconate-specific defect (**Figure S2C**). The *rafB* and *ogl/kdgM* mutants showed relatively mild defects under aerobic conditions, but these became markedly more pronounced under anaerobic conditions, which better reflect the intestinal environment (**Figures 2F and G**). In these conditions, such mutants also exhibited a selective disadvantage in competing with WT MH258 only when grown on the corresponding carbon source, but not on glucose or on a related parent sugar (sucrose for *rafB*) (**Figure S4A**). Importantly, the fitness defect of each individual knockout was retained when competition assays were performed in medium containing a mixture of five carbohydrates (raffinose, arabinose, arabitol, xylose, and gluconate) indicating that each of these sugars continues to contribute to *K. pneumoniae* fitness even when multiple alternative carbon sources are simultaneously available (**Figures S5A–G**).

We next evaluated the contribution of these pathways to intestinal colonization *in vivo*. Antibiotic-treated mice (AMNV, 4 days) were gavaged with a 1:1 mixture of WT and mutant MH258, and relative fitness was assessed over time by competitive index analysis using selective plating of fecal samples (**Figure 2H**). These experiments revealed that most of the tested carbon utilization pathways are required for efficient colonization of the antibiotic-treated intestine, at least in this mouse model (**Figures 2I-O**). In particular, mutants defective in arabinose, raffinose and xylose utilization (*araA*, *rafB* and *xylA*) exhibited strong competitive defects, as did the *gntK* mutant, in line with previous data ^28^, although the latter mutation may have pleiotropic metabolic effects. Consistently, the *gntT/U* transporter mutant displayed a much milder *in vivo* defect (**Figure 2I**). For selected mutants displaying an *in vivo* fitness defect, plasmid-based complementation restored growth on the corresponding carbon source *in vitro* (**Figures 2B–G and S4B–C).** Except for the oligogalacturonate mutant, all *in vivo* phenotypes could be rescued by complementation (**Figures 2I–O**). For the *kdgM/ogl* double knockout, *in vivo* complementation was unsuccessful (**Figure 2G**), likely due to two factors: (i) deletion of *kdgM/ogl* resulted in downregulation of additional oligogalacturonate genes within the same genomic island, whose expression did not return to baseline upon complementation (**Figure S4C**); and (ii) the complementation plasmid was lost *in vivo* in at least 50% of bacteria, as assessed by PCR (data not shown). Deletion of genes involved in sorbitol (*srlA*), ethanolamine (*eutE*), and glucuronate (*uxaC*) utilization had little or no impact on colonization (**Figure 2I**). Together, these data demonstrate that transcriptomic profiling reliably identifies carbon sources that are functionally important for *Kp* colonization of the antibiotic-disrupted intestine. Notably, several of the pathways uncovered here, including arabinose, raffinose, arabitol, and oligogalacturonate utilization, have not previously been implicated in *Kp* intestinal colonization, while others, such as gluconate and xylose metabolism, confirm and refine prior observations in a microbiota-depleted model that is directly relevant to the human setting.

### Impacts of dietary changes on *K. pneumoniae* carbon source utilization

Dietary composition profoundly influences the biology of intestinal microbes, not only by favoring specific strains but also by shaping the metabolic strategies available to each bacterium, as it determines which carbon sources are present in the gut ^34,41–44^. Our initial experiments were performed in mice fed a standard rodent maintenance diet (STD, SAFE150), which is cereal-based, rich in plant-derived carbohydrates, and relatively low in fats (**Figure S6A**). In contrast, Western diets are typically enriched in animal-derived fats and proteins, as well as in refined sugars. To determine whether our findings were dependent on the diet used or instead reflected more general principles of nutrient utilization in the intestine, we repeated the colonization and transcriptional analyses in mice fed a high-fat diet containing simple sugars (sucrose) and fats derived from butter and animal-derived proteins (HFD, SAFEU8954 supplemented with 1.25% cholesterol) (**Figures 3A and S6A**). *Kp* robustly colonized mice fed either diet, although bacterial densities were consistently lower under HFD than under STD conditions. This quantitative difference was reproducibly observed *in vivo*, in *ex vivo* assays using cecal contents from antibiotic-treated mice, and *in vitro* in water-resuspended chow (**Figure S6B**). This suggests that, despite its higher caloric content, the HFD provides a less favorable nutritional environment for *Kp*. Consistent with this interpretation, metabolomic analysis of cecal contents from antibiotic-treated mice showed that HFD feeding markedly reduced the abundance of raffinose, arabinose, arabitol, xylose, and gluconate, all of which had been identified as carbon sources supporting *Kp* intestinal fitness (**Figure S7A and and S7C-L**).

**Figure 3.**
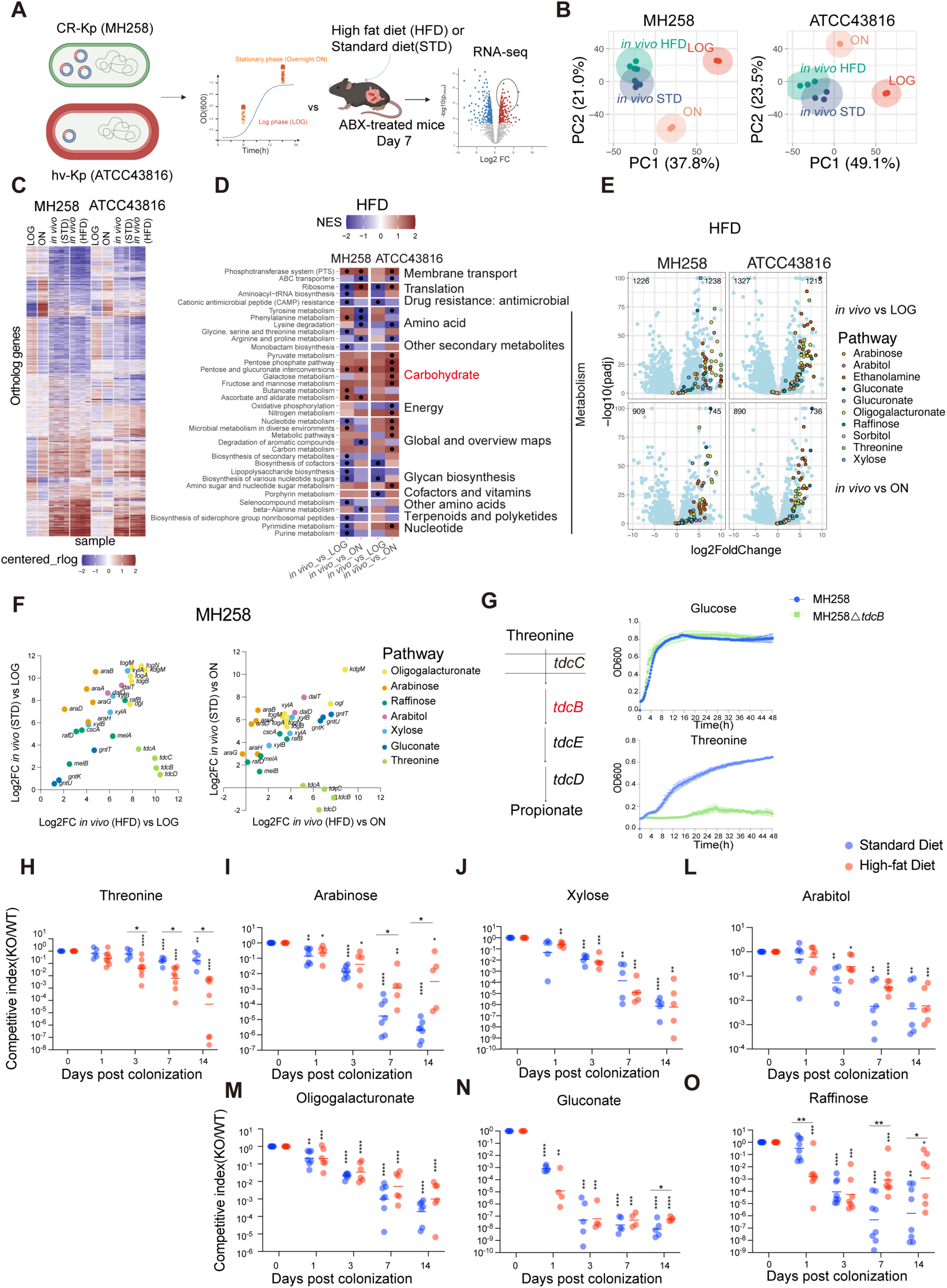
Dietary composition modulates carbon utilization strategies during *K. pneumoniae* intestinal colonization. **(A)** Experimental schematic illustrating transcriptomic analysis of *K. pneumoniae* colonizing antibiotic-treated mice fed either a standard diet (STD) or a high-fat diet (HFD). Data from STD-fed mice are also shown in Figure 1. **(B)** Principal component analysis (PCA) of *K. pneumoniae* transcriptomes from mice fed a standard diet (STD) or a high-fat diet (HFD), together with *in vitro* controls. Each dot represents one sample. **(C)** Heatmap and hierarchical clustering of global gene expression profiles from *in vivo* samples collected from STD- and HFD-fed mice. Columns represent individual samples and rows represent genes. Relative gene expression is shown, with red indicating upregulation and blue indicating downregulation compared to *in vitro* conditions. **(D)** KEGG pathway enrichment analysis of genes differentially expressed *in vivo* relative to *in vitro* in both strains under HFD conditions. Red indicates pathways enriched among upregulated genes. Black dots denote significantly enriched pathways. **(E)** Volcano plots displaying differential gene expression of *K. pneumoniae in vivo* relative to *in vitro* for both strains. Light blue dots indicate significantly differentially expressed genes (|log2FC| > 1, padj < 0.05), whereas gray dots indicate non-significant genes. Metabolic pathways/carbon sources of interest are color-coded as in Figure 1e, with the addition of threonine-associated genes specifically observed in HFD-fed mice. **(F)** Correlation analysis of differential expression of genes associated with seven key carbon sources in *K. pneumoniae* (MH258) colonizing mice fed STD versus HFD. Each dot represents one gene, with colors indicating different carbon utilization pathways. **(G)** Threonine utilization was disrupted by deletion of *tdcB*, and WT and KO strains were grown in M9 medium supplemented with threonine as the sole carbon source. Shown is one of three independent experiments. **(H-O)** Competitive fitness of the indicated mutant and complemented mutant strains relative to WT in mice maintained on a standard diet or high-fat diet, as indicated. Competitive indices were normalized to the corresponding inoculum ratio and are displayed on a logarithmic scale. A competitive index of 1 indicates no change relative to the inoculum. Each point represents one mouse, data are pooled from 2–3 independent experiments, and horizontal lines indicate the geometric mean. Statistical analyses were performed on log10-transformed competitive-index values. Symbols above individual groups indicate significance relative to a competitive index of 1, whereas brackets indicate direct comparisons between groups. Comparisons with the neutral value were performed using two-sided one-sample t-tests, and direct comparisons between dietary conditions were performed using two-sided unpaired Welch’s t-tests (\**P* < 0.05; \*\**P* < 0.01; \*\*\**P* < 0.001; \*\*\*\**P* < 0.0001).

RNA sequencing revealed that the overall *in vivo* transcriptional programs of both MH258 and ATCC 43816 were largely conserved across the two diets and remained clearly distinct from those observed *in vitro* (**Figures 3B and 3C, and S6C and D**). In both diets, the most prominent transcriptional alterations involved metabolic pathways, particularly those associated with carbohydrate utilization (**Figure 3D**). At the gene level, the metabolic pathways previously identified under STD conditions were also induced in HFD-fed mice relative to *in vitro* growth. These included pathways involved in the utilization of arabinose, arabitol, gluconate, oligogalacturonate, raffinose, and xylose, indicating that these substrates continue to contribute to *Kp* metabolism under HFD conditions (**Figure 3E**). However, the magnitude of induction differed between diets. In particular, genes involved in arabinose and oligogalacturonate utilization were expressed at lower levels under HFD than under STD conditions in both MH258 and ATCC43816. Conversely, threonine catabolism genes were significantly induced relative to *in vitro* conditions only in HFD-fed mice and were therefore also more highly expressed under HFD than under STD conditions (**Figures 3F and Figure S6C-E**). These findings indicate that, although the overall metabolic program remained largely conserved, HFD reshaped the relative contribution of individual nutrients by reducing reliance on some carbohydrate sources while increasing threonine utilization. Metabolomic analyses supported this interpretation. Although threonine abundance was comparable in the cecal contents of uncolonized STD- and HFD-fed mice, MH258 colonization significantly depleted threonine only under HFD conditions, consistent with its increased utilization in this dietary context (**Figure S7B**). Conversely, raffinose, arabinose, arabitol, xylose, and gluconate were substantially less abundant in HFD-fed than in STD-fed mice but were nevertheless significantly depleted following MH258 colonization under HFD conditions (**Figures S7A and S7C-L**). Thus, *Kp* continues to consume the carbon sources identified under STD conditions despite their reduced availability in the HFD-fed intestine, while simultaneously increasing its reliance on threonine.

We next evaluated the fitness of MH258 metabolic mutants characterized above in HFD-fed mice using competitive index experiments, including an additional *tdcB* (threonine dehydratase) mutant validated *in vitro* (**Figure 3G**). Notably, all tested mutants exhibited fitness defects in HFD-fed mice (**Figures 3H-O**). Mutants impaired in arabitol, xylose, oligogalacturonate, and gluconate utilization showed phenotypes comparable to those observed under the standard diet. By contrast, the *araA* and *rafB* mutants showed weaker defects under HFD, whereas the *tdcB* mutant exhibited a more pronounced defect, consistent with the diet-dependent changes predicted by the transcriptomic and metabolomic analyses (**Figures 3H-O and S6E**).

Collectively, these results show that *Kp* retains a core set of nutrient requirements across distinct diets while adjusting its reliance on individual substrates according to dietary context. They further demonstrate that *in vivo* transcriptional profiling can identify diet-specific metabolic requirements, while integration with metabolomic data helps define which nutrient dependencies are most relevant in a given nutritional environment.

### *K. pneumoniae* carbon source utilization may be conserved in the human intestine

The above results prompted us to ask whether *Kp* utilizes similar carbon sources during colonization of the human intestine. To this end, we analyzed stool samples from three patients colonized with *Kp*, one of whom provided two samples two weeks apart (**Figure 4A and Table S1**). Levels of *Kp* colonization varied substantially among samples, ranging from 3% to 73%, and so did the composition of the background microbiota (**Figure 4B**). Two of the colonizing isolates, corresponding to three samples, belonged to sequence type ST437 (clonal group 258, as is strain MH258), whereas one isolate belonged to ST1427, a sequence type not typically associated with multidrug resistance (**Figure 4B**).

**Figure 4.**
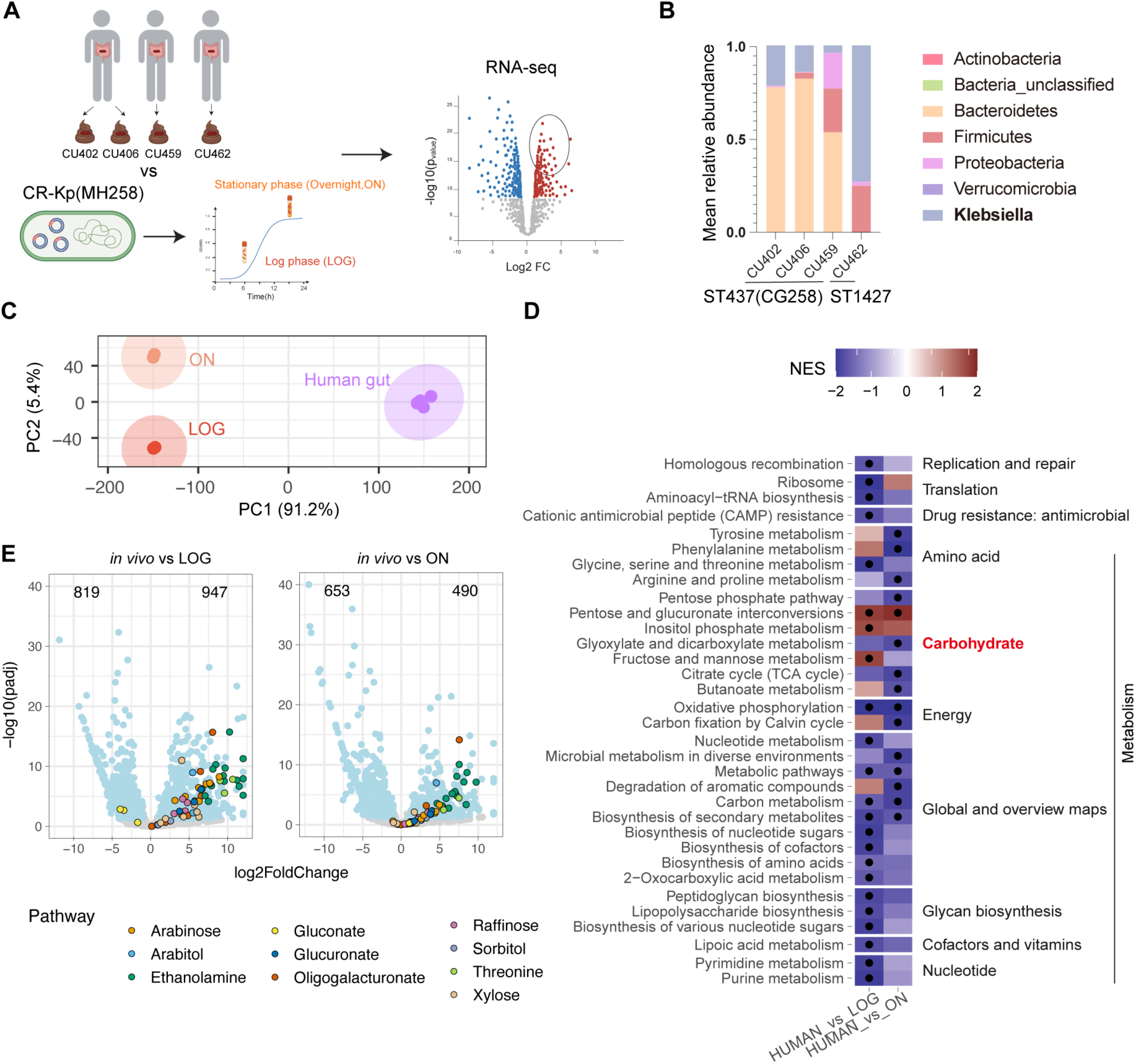
*K. pneumoniae* carbon utilization signatures are conserved in the human gut. **(A)** Experimental workflow. Four fecal samples were collected from three antibiotic-treated patients colonized with *K. pneumoniae*, including two samples obtained from the same patient at different treatment time points. Bacterial RNA was extracted and subjected to RNA sequencing. Transcriptomes obtained from these samples were compared with that from *in vitro*–grown MH258 (data shown in Figure 1) to identify genes upregulated during intestinal colonization. See also Table S1 for additional information on patients. **(B)** Microbiota composition and *K. pneumoniae* abundance in patient fecal samples determined by 16S rRNA gene sequencing. WGS (Whole-genome sequencing)-based sequence typing of *Klebsiella* is also shown. **(C)** Principal component analysis (PCA) of *K. pneumoniae* transcriptomes derived from human fecal samples and *in vitro* cultures. Each dot represents one sample. **(D)** KEGG pathway enrichment analysis of genes significantly upregulated in the human gut relative to *in vitro* culture. **(E)** Volcano plots depicting differential gene expression of *K. pneumoniae* in the human gut relative to *in vitro* conditions. Each dot represents one gene. Light blue dots indicate significantly differentially expressed genes, whereas gray dots indicate non-significant genes. Genes associated with carbon sources identified in the mouse model are highlighted using the color code adopted in Figures 1E and 3E.

We extracted bacterial RNA from these samples and compared *Kp* gene expression profiles to those obtained from *in vitro*–grown bacteria, using MH258 as the reference and restricting the analysis to genes shared across strains (see Methods for details). Principal component analysis revealed a clear separation between *in vivo* and *in vitro* transcriptomes (**Figure 4C**), accompanied by broad changes in pathway regulation (**Figure 4D**). Notably, many of the metabolic genes identified as upregulated *in vivo* in the mouse model across dietary conditions were similarly upregulated in human samples (**Figure 4E**). An exception was the gluconate utilization pathway, which appeared downregulated *in vivo* relative to logarithmic-phase *in vitro* cultures (**Figure 4E**). Because strain-matched *in vitro* controls were not used, this analysis cannot establish the precise magnitude of transcriptional regulation for each individual isolate. Nevertheless, the recurrence of a similar metabolic signature across genetically distinct human isolates, despite differences in strain background, residual microbiota composition, and diet, argues that these transcriptional changes are not specific to MH258 and supports the existence of shared nutrient-utilization programs during intestinal colonization in mice and humans. Together, these findings suggest that the carbon sources identified in this study may also be relevant during human intestinal colonization and warrant further investigation as potential targets for therapeutic intervention.

### Rational optimization of metabolic competitors for *K. pneumoniae*

Having identified key metabolic determinants of *Kp* gut colonization in mice, and putatively in humans, we next explored whether these pathways could be targeted through metabolic interference using rationally assembled probiotics. Indeed, nutrient competition–based displacement represents a promising approach to decolonize intestinal pathobionts, including *Klebsiella* ^26,28,36,37,45–47^. However, because no single microbial competitor fully overlaps with the metabolic niche of *Kp*, effective decolonization is thought to require consortia of multiple microbes, both in preventive and therapeutic settings ^26,28,36,37^. Despite the broad carbon utilization capacity of *Enterobacteriaceae* ^31^, a sufficient degree of baseline niche overlap within this family enables partial competitive exclusion ^26^. For example, *Klebsiella oxytoca* can outcompete *Klebsiella pneumoniae* over multiple carbon sources ^36^, and *Klebsiella sp.* ARO112 can suppress pathogenic *Escherichia coli* ^45,^*^4C^*, while conversely *E. coli* has been shown to enhance colonization resistance against *Kp* when incorporated in commensal microbial consortia ^26^. Based on these observations, we reasoned that rationally enhancing carbon source utilization in a competitive *Enterobacteriaceae* could improve colonization resistance against *Kp*.

We selected *E. coli* Nissle 1917 (*EcN*) as a chassis, as it is commercially available as a probiotic, considered safe even in hosts with compromised intestinal barriers, and fully genetically tractable ^48,49^. Furthermore, this strain, in combination with another *E. coli*, could provide colonization resistance against pathogenic *Enterobacteriaceae* ^50^. In mouse colonization experiments, *EcN* exhibited a carbohydrate utilization profile largely overlapping with that of MH258 (**Figure 5A**), although notable exceptions were observed. For instance, *EcN* was unable to consume arabitol, consistent with the absence of the corresponding metabolic genes (**Figure 5B**). Raffinose utilization was also incomplete: in *Klebsiella*, raffinose is hydrolyzed by MelA into sucrose and galactose, with sucrose further cleaved into glucose and fructose by CscA (**Figure 5B**). As *EcN* lacks *cscA*, raffinose breakdown was partial, resulting in lower raffinose levels than in mock-reconstituted controls, but higher sucrose levels compared to *Kp*-colonized mice.

**Figure 5.**
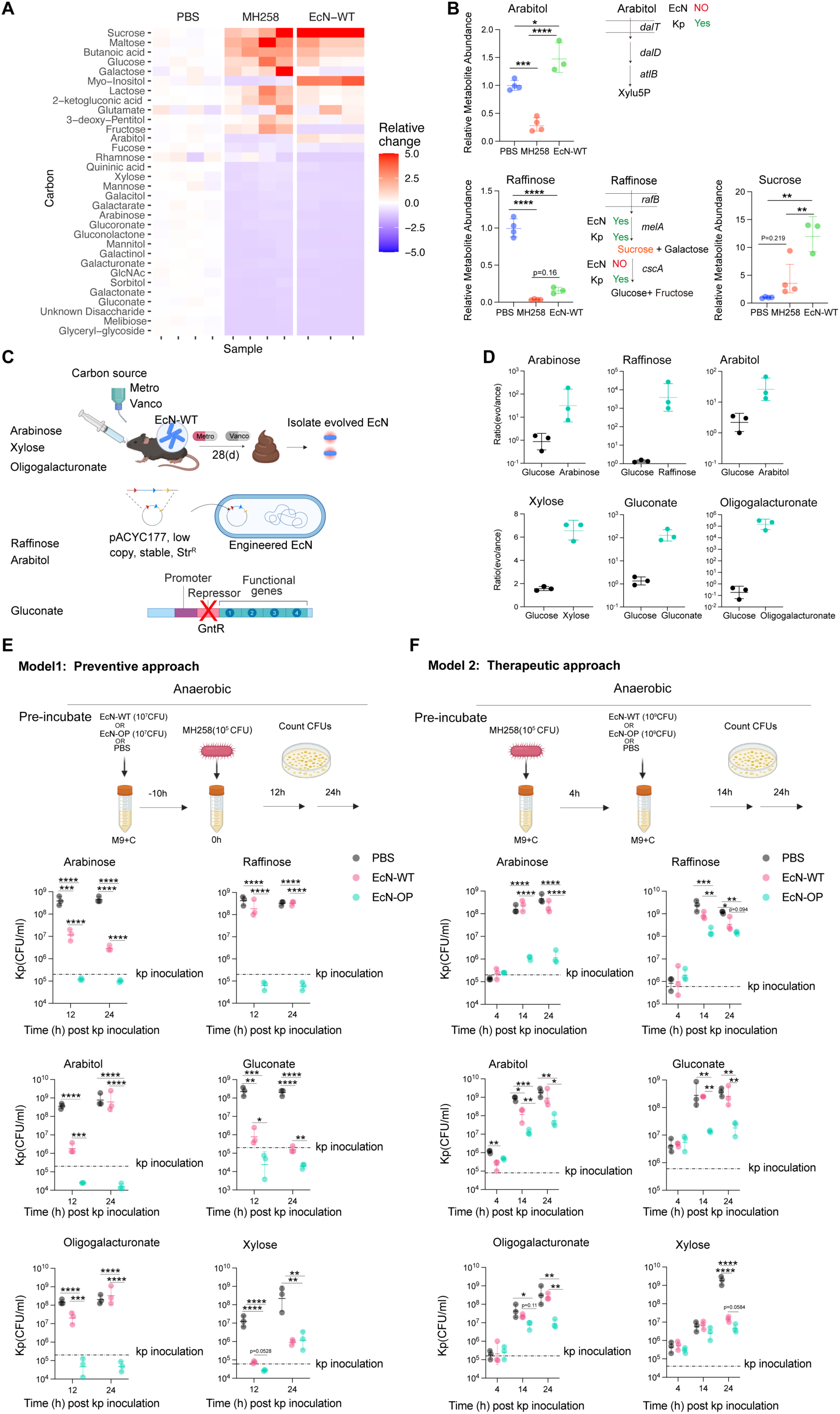
Optimization of carbon utilization pathways in *Escherichia coli* Nissle enhances competition with *K. pneumoniae in vitro*. **(A)** GC–MS–based metabolomic profiling of carbon source utilization by *E. coli* Nissle (EcN-WT) during intestinal colonization, analyzed using the same approach displayed in Figure 1. Heatmap shows relative changes in metabolite abundance compared with PBS-treated controls. Red indicates increased abundance, blue indicates decreased abundance, and white indicates no substantial change in any given carbon source. Columns represent individual mice (n=4 for PBS and MH258, n=3 for EcN-WT). **(B)** GC–MS or LC–MS quantification of the depicted sugars in the intestine of mice colonized with MH258 or EcN-WT; the presence/absence of metabolizing enzymes in the respective genomes is also shown. Each dot represents one mouse; horizontal lines indicate geometric means ± geometric SD. **(C)** Schematic overview of the three strategies used to optimize EcN for improved utilization of key carbon sources. For arabinose, xylose, and oligogalacturonate, *in vivo* experimental evolution was performed in mice supplied with 0.6% of the indicated carbon source in drinking water (see Figure S6A for more details). For raffinose and arabitol, relevant metabolic genes from *K. pneumoniae* were expressed in EcN-WT using a low-copy, stable plasmid (pACYC177). For gluconate, deletion of the transcriptional repressor GntR was used to enhance expression of gluconate utilization genes. The resulting strains are referred to as Optimized *E. coli* Nissle (EcN-OP). **(D)** *In vitro* anaerobic competition assays comparing carbon source utilization between EcN-OP and EcN-WT. Strains were co-cultured at a 1:1 ratio in M9 minimal medium supplemented with glucose or individual carbon sources of interest. EcN-OP refers to the respective engineered/evolved strain (e.g., arabinose-evolved strain was tested in M9 medium supplemented with glucose or arabinose). Shown is one representative of three independent experiments. **(E)** *In vitro* protection model assessing the ability of EcN-OP to prevent *K. pneumoniae* expansion. Minimal M9 medium containing glucose or individual carbon sources was inoculated with PBS, EcN-WT or the respective EcN-OP (10^7^CFU). 10 hours later, MH258 was also inoculated (10^5^CFU), and bacterial abundance was quantified by selective plating at the indicated time points. Shown is one representative of three independent experiments. **(F)** *In vitro* decolonization model assessing the ability of EcN-OP to suppress *K. pneumoniae* following prior establishment. MH258 was first inoculated (10^5^ CFU) and allowed to establish in minimal medium containing glucose or individual carbon sources, after which EcN-OP, EcN-WT (10^9^ CFU), or PBS were introduced. Bacterial abundance was quantified at the indicated time points by selective plating. Shown is one representative of three independent experiments. For all experiments, statistical significance was determined using one-way ANOVA followed by Tukey’s multiple comparisons test (\**P* < 0.05; \*\**P* < 0.01; \*\*\**P* < 0.001; \*\*\*\**P* < 0.0001). In (E) and (F), dashed lines indicate the initial inoculum level of *K. pneumoniae*.

To enhance utilization of the identified carbon sources by *EcN*, we explored several strategies. First, we attempted to increase expression of relevant operons by replacing native promoters with a strong constitutive promoter (**Figure S8A**). However, for both gluconate and arabinose metabolic operons, this approach resulted in reduced bacterial growth despite robust upregulation of target genes (**Figures S8B-G**), indicating that indiscriminate overexpression can be detrimental for bacterial fitness. We therefore pursued three alternative strategies: (i) for carbon sources already accepted by *EcN*, we performed *in vivo* adaptive evolution in mono-colonized mice exposed to specific carbon sources in drinking water; (ii) in case of metabolic genes absent in *EcN*, we opted for heterologous expression of *Klebsiella* genes; and (iii) for gluconate, whose genetic locus organization allowed for this, we deleted the regulator *gntR*, derepressing transcription of the controlled metabolic genes (**Figure 5C**). Methodological details and characterization of gene expression and growth kinetics for engineered and evolved strains are provided in **Figures S9A–U**. All optimized *EcN* strains (*EcN*-OP) exhibited enhanced growth and fitness relative to the parental strain when cultured in the corresponding carbon source, but not in glucose, indicating improved substrate-specific metabolic capacity (**Figures 5D and S9 C-U**). To identify the genetic changes underlying the evolved phenotypes, we performed whole-genome sequencing of *EcN*-OGA and *EcN*-Ara, which exhibited the strongest fitness gains among the adaptively evolved strains. *EcN*-OGA carried a loss-of-function mutation in *kdgR*, which encodes a repressor of oligogalacturonate utilization genes, whereas *EcN*-Ara harbored a missense mutation in *araC*, the transcriptional regulator of the arabinose utilization pathway. These mutations likely account for the enhanced ability of the respective strains to utilize their corresponding carbon sources **(Figure S9V)**.

We next assessed whether these optimized strains exhibited superior competitive ability against *Kp in vitro*. Using two complementary experimental setups, modeling protection (*EcN* added before *Kp*) and treatment (*EcN* added after *Kp*) respectively, similar to what described previously ^26^, we evaluated metabolic displacement in media containing the relevant carbon sources. In all cases, optimized *EcN* strains outperformed the WT strain, resulting in increased relative *EcN* abundance and more efficient displacement of *Kp* for both MH258 and ATCC 43816 (**Figures 5E and F, S10A and B, and S11A-F**). Collectively, by combining *in vivo* evolution and targeted genetic engineering, we generated six optimized *EcN* strains with enhanced potential to metabolically displace *Kp* through consumption of raffinose, arabinose, arabitol, gluconate, oligogalacturonate, and xylose.

### A rationally assembled consortium of optimized *E. coli* Nissle strains provides enhanced colonization resistance against *K. pneumoniae*

We next assembled a consortium composed of the six optimized *E. coli* Nissle strains generated above, combined in equal proportions, to test their efficacy *in vivo*. This consortium was termed NICHE (*Nissle Integrated Consortium for Heterogeneous Energy utilization*) (**Figure 6A**). Tracking of the genetically barcoded strains following gavage into antibiotic-treated mice revealed progressive shifts in consortium composition, with *EcN*-Raf dominating by day 14 and *EcN*-Gnt and *EcN*-Xyl declining to low abundance (**Figures S12A–D**). At day 7, NICHE depleted raffinose and arabitol relative to *EcN*-WT, whereas changes in arabinose, xylose, and gluconate were not visible, potentially because effects occurred earlier when the corresponding strains were more abundant; oligogalacturonate could not be quantified due to technical limitations (**Figures S12E–L**). Because nutrient availability and strain fitness may vary over time and across hosts, we retained all six strains to maximize metabolic coverage in subsequent experiments.

**Figure 6.**
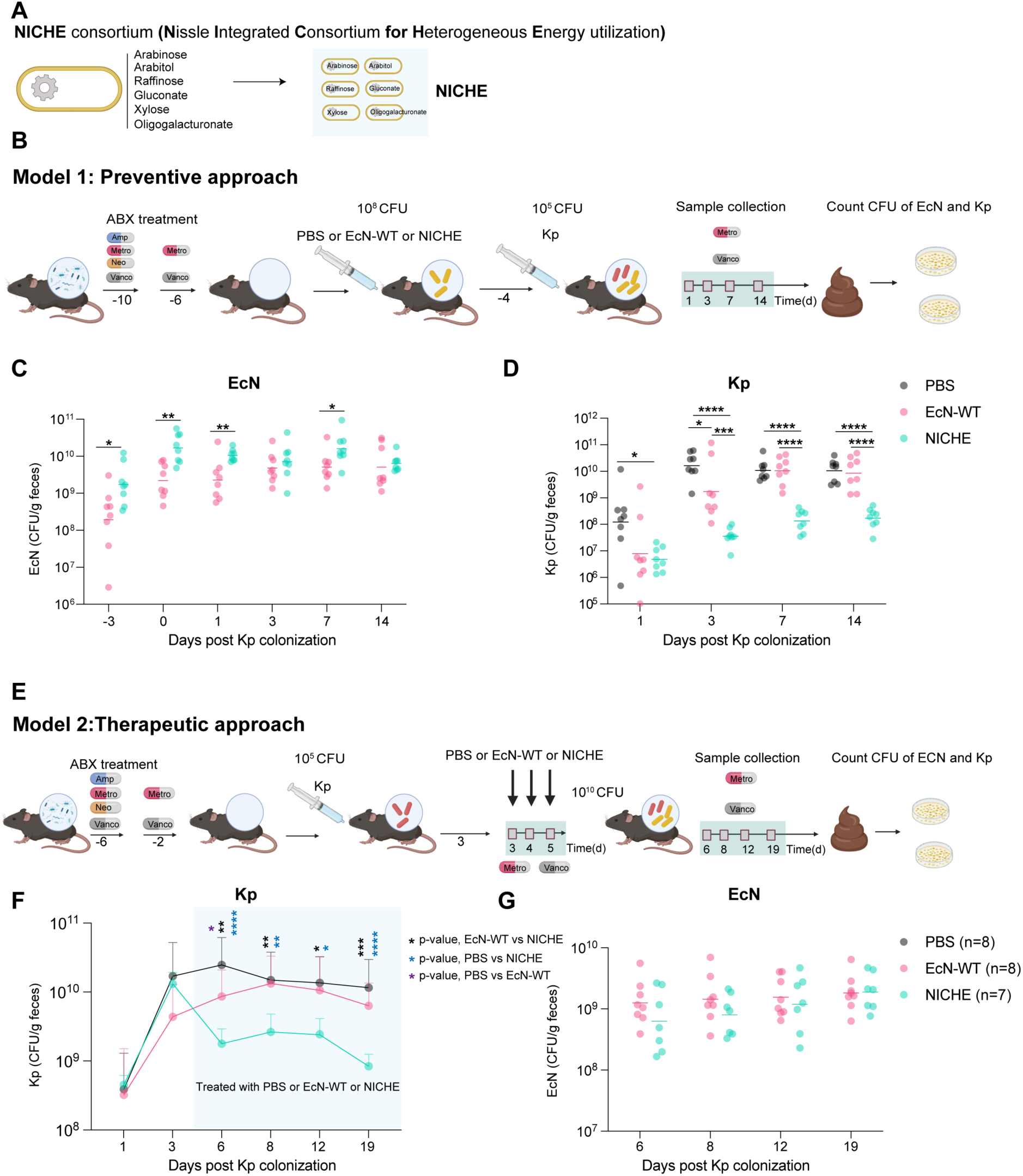
An optimized *E. coli* Nissle consortium suppresses *K. pneumoniae* intestinal colonization *in vivo*. **(A)** Six EcN-OP strains optimized for utilization of distinct carbon sources (arabinose, arabitol, raffinose, oligogalacturonate, gluconate, and xylose) were assembled into a defined consortium (NICHE). **(B)** To evaluate protection, mice were treated with AMNV for 4 days, followed by a flush-out of 2 days in MV, then pre-colonized with NICHE, EcN-WT, or administered an equal volume of PBS by single gavage. After 4 days, mice were challenged with *K. pneumoniae* (10^5^ CFU). Fecal bacterial burdens were monitored longitudinally. **(C)** Fecal density of EcN-WT or NICHE in colonized mice at the indicated time points. **(D)** Fecal burden of *K. pneumoniae* MH258 at the indicated time points from mice previously reconstituted with PBS, EcN-WT or NICHE. Shown are individual mice and geometric means. n=8, data pooled from 2 independent experiments. **(E)** To evaluate decolonization, antibiotic-treated mice (same regimen as in (B)) were first colonized with *K. pneumoniae*. After 3 days, mice were administered NICHE, EcN-WT, or PBS by gavage on three subsequent days. **(F)** Fecal burden of *K. pneumoniae* MH258 at the indicated time points mice depcted in (E). Shown are individual mice and geometric means. n=7-8, pooled from two independent experiments. **(G)** Density of EcN-WT or NICHE in the fecal pellets of mice shown in (E) and (F). Shown are individual mice and geometric means. n=4 from one representative of two independent experiments. Statistical significance was determined using unpaired two-tailed t tests for comparisons between two groups and one-way ANOVA followed by Tukey’s multiple comparisons test for comparisons involving more than two groups (\**P* < 0.05; \*\**P* < 0.01; \*\*\**P* < 0.001; \*\*\*\**P* < 0.0001).

We first tested NICHE prophylactically in mice maintained on metronidazole and vancomycin after four days of AMNV treatment. Animals received EcN-WT or NICHE (10⁸ CFU), were challenged four days later with MH258 (10⁵ CFU), and were monitored longitudinally by selective plating of fecal samples (**Figure 6B**). Both *EcN* WT and NICHE rapidly expanded in the gut during the first four days after gavage, a period during which NICHE showed a modest but reproducible growth advantage. From day 7 post–*EcN* gavage (day 3 post–*Kp* challenge), the densities of the two *E. coli* populations converged at approximately 10¹⁰ CFU/g feces and remained stable thereafter (**Figure 6C**). Strikingly, at all time points analyzed, NICHE, but not *EcN* WT, conferred significant protection against *Kp*, which expanded to markedly lower levels in NICHE-colonized mice. In particular, between 3 and 14 days post infection, NICHE reduced *Kp* burdens by 2–3 orders of magnitude compared with PBS- or *EcN* WT–treated controls (**Figure 6D**). Importantly, supplementation of the six target carbohydrates in the drinking water markedly attenuated the protective effect of NICHE, restoring *Kp* colonization to levels comparable to those in PBS-treated mice by day 14 (**Figure S13D**), supporting nutrient competition as the primary mechanism underlying NICHE-mediated colonization resistance. Because *EcN*-Raf and *EcN*-OGA dominated *in vivo* and NICHE preferentially depleted raffinose and arabitol, we asked whether protection required the full consortium or was driven mainly by strains targeting the most relevant substrates. A simplified consortium (mini-NICHE), comprising the raffinose-, arabitol-, and oligogalacturonate-optimized strains (**Figure S13A**), showed efficacy comparable to complete NICHE *in vivo* (**Figures S13E–F**). This indicates unequal strain contributions to protection, while also showing that retaining additional engineered strains is not detrimental and may broaden efficacy when dietary carbon-source availability is variable or unknown.

To assess whether NICHE also limits the pathogenic potential of hypervirulent *Kp*, we examined its impact on intestinal colonization and bacterial dissemination by the hv-*Kp* strain ATCC 43816. Mice colonized with *EcN* WT or NICHE were challenged orally with ATCC 43816 (10^4^ CFU), and bacterial burdens were quantified one and three days later (**Figure S14A**). As observed for MH258, NICHE significantly restricted intestinal expansion of ATCC 43816 compared with PBS- or *EcN* WT–treated controls, despite comparable *EcN* WT and NICHE densities (**Figures S14B and C**). Gentamicin treatment of the ceca eliminated this difference, indicating that NICHE primarily reduced bacteria associated with the cecal surface rather than those that had invaded the tissue (**Figures S14D and E**). Importantly, bacterial burdens in the mesenteric lymph nodes, liver, and spleen tended to be lower in NICHE-reconstituted mice, although the differences were not statistically significant, and were undetectable in several animals. Accordingly, a higher proportion of mice in this group had no detectable CFU in the extraintestinal organs based on the corresponding limits of detection, suggesting that NICHE may help limit systemic dissemination of hypervirulent *Kp* (**Figures S14F–H**).

Under HFD conditions, the abundance of several carbon sources targeted by NICHE, including raffinose and arabinose, was reduced (Figure S7). Consistent with this altered nutrient landscape, the *in vivo* fitness defects associated with disruption of the corresponding *K. pneumoniae* utilization pathways were also diminished under HFD (Figures 3H–O). HFD itself constrained *K. pneumoniae* expansion, and both EcN-WT and NICHE further reduced colonization relative to HFD-fed controls (Figures S14I–K). However, NICHE did not provide a detectable additional benefit over EcN-WT under HFD, in contrast to the stronger protection provided by NICHE under the standard-diet conditions tested here. Together, these findings indicate that the additional benefit conferred by the engineered metabolic functions is dependent on dietary nutrient availability and diminishes when the targeted substrates become less abundant.

We next tested a therapeutic (decolonization) model, in which mice were first administered *Kp* MH258, mimicking the clinical scenario in which an antibiotic-treated patient becomes colonized. Starting three days later, mice were gavaged with *EcN* WT or NICHE (10^10^ CFU) for three consecutive days (**Figure 6E**). Once again, NICHE treatment resulted in a significant reduction of *Kp* burdens compared with both mock treatment and *EcN* WT, despite comparable intestinal expansion of the two *E. coli* populations (**Figures 6F and 6G**). Together, these data demonstrate that treatment with a rationally engineered microbial consortium targeting transcriptomics-identified carbon sources provides protection against *K. pneumoniae* intestinal colonization in both preventive and therapeutic settings, with the magnitude of this benefit shaped by dietary nutrient availability.

## Discussion

Intestinal colonization is a critical step in the pathogenesis of antibiotic-resistant *K. pneumoniae* infections. Colonization resistance is largely governed by competition for limiting resources ^51^, with organisms that gain superior nutrient access acquiring a fitness advantage ^25,27^. As a gut-adapted pathobiont, *Klebsiella* can persist at low abundance within complex microbial communities yet rapidly expand following antibiotic exposure, frequently reaching domination levels that strongly predict subsequent invasive infection ^5,8,9^. By depleting commensal competitors, antibiotics release nutrients such as monosaccharides and sugar acids ^20^, creating transient nutrient niches that *Klebsiella* is well positioned to exploit ^31^. Here, we developed a framework to identify these limiting carbon sources and leverage it to reinforce colonization resistance.

We inferred intestinal carbon-source utilization by comparing *Kp* transcriptomes *in vitro* and *in vivo*. Although related strategies have been reported, our results show that this approach is robust across strains and diets, with high sensitivity and specificity. Transcriptomics sensitively revealed pathways engaged *in vivo*, although substrates used similarly *in vitro* and *in vivo* may escape differential-expression analysis, while pathways linked to scarce intestinal nutrients may appear disproportionately important. Metabolomics therefore provided an essential complement by confirming consumption and measuring relative substrate availability, which emerged as an important determinant of nutrient contribution to fitness.

In the antibiotic-treated intestine, *Kp* did not primarily rely on broadly accessible substrates such as glucose, but instead exploited a restricted set of carbon sources, including arabinose, arabitol, raffinose, and (oligo)galacturonate. Other sugars, such as fucose, previously implicated in intestinal colonization ^30^, were also supported by our transcriptomic and metabolomic datasets. Two unrelated strains from distinct clonal groups and pathotypes displayed similar preferences, consistent with the near-universal conservation of the corresponding utilization genes across *Klebsiella* genomes ^31,32^. Other nutrients, such as gluconate ^28^ appeared more context dependent, potentially reflecting strain-specific metabolic wiring ^31,32^ or differences in availability. Thus, post-antibiotic expansion appears to rely on efficient exploitation of a limited number of conditionally available niches.

Although utilization of the principal carbon sources was largely preserved across standard and high-fat, high-sugar diets, diet altered the metabolic context in which these pathways operated. Under high-fat-diet conditions, depletion of otherwise preferred substrates was accompanied by a selective contribution of threonine utilization, suggesting increased dependence on secondary nutrients when primary substrates are scarce.

Diet also influenced absolute pathogen expansion, highlighting that a pathway’s contribution to relative fitness does not necessarily predict its effect on total burden. Importantly, competitive defects measured between isogenic wild-type and utilization-deficient strains may overestimate the impact of restricting a single nutrient when *Kp* competes with metabolically distinct organisms, as differences in their broader metabolic repertoires can provide alternative routes for nutrient exploitation. A substrate may confer a competitive advantage without being abundant enough to sustain extensive growth, whereas diets that restrict preferred nutrients may lower carrying capacity while exposing narrower residual dependencies amenable to probiotic targeting. Host-derived glycans^52^ and microbial cross-feeding ^53^, further shape this nutrient landscape.

Based on the acquired knowledge, we next sought to reinforce colonization resistance using a rationally engineered probiotic. Competitive exclusion is driven by nutrient overlap ^25–27^. However, the broad metabolic flexibility of *Kp* limits the suppressive capacity of individual commensals. ^31,32^ Defined consortia may be more effective by collectively consuming multiple limiting substrates ^28,36,37^ and organisms such as *E. coli* may therefore contribute substantially to colonization resistance through broader-community “higher-order effects” ^26^.

Our findings extend this framework by showing that metabolic overlap can be enhanced within a single probiotic chassis. Using *E. coli* Nissle, we combined genetic engineering and *in vivo* evolution to increase consumption of carbon sources preferred by *Kp*. The resulting NICHE consortium imposed metabolic interference, reduced the pathogen’s fitness ceiling, and prevented domination. Although NICHE did not eliminate *Kp*, partial suppression may suffice to restore ecological balance and reduce infection risk. Human studies suggest that systemic infection risk decreases markedly below approximately 20% relative abundance ^9,54^, indicating that reductions of one to two orders of magnitude may be protective. Moreover, NICHE remained effective under continuous antibiotic exposure, despite persistent microbiota disruption and continued generation of niches favoring pathogen expansion.

The unequal persistence of individual NICHE strains, together with the comparable efficacy of the simplified consortium, indicates that not all members need to be stably maintained for activity. Although simplification may improve manufacturability of the consortium, broader diversity may remain advantageous when nutrient availability is unpredictable, as it does not appear to impair consortium fitness and may increase flexibility across carbon-source environments.

To further enhance the potential of engineered consortia, probiotics could be coupled with interventions that reshape the ecological environment in their favor. The additional benefit provided by the engineered functions was dependent on dietary context. Under the standard-diet conditions tested here, where the targeted substrates were more abundant, NICHE provided greater protection than parental EcN. In contrast, under HFD, where several targeted nutrients were reduced and the corresponding *K. pneumoniae* utilization pathways made smaller contributions to fitness, the additional benefit of NICHE over EcN-WT was diminished. This diet dependence represents an important boundary condition of targeted nutrient competition: the benefit conferred by an engineered metabolic function depends on the availability of its target substrate. Accordingly, maximal efficacy may require matching the metabolic capabilities of an engineered consortium to the nutrient landscape associated with a particular dietary or host context. Probiotics could also be combined with dietary strategies that increase antibiotic sensitivity ^55,56^ or reduce virulence factor expression ^57^, as well as with vaccine-induced immunity ^58^, engineered antimicrobial production ^59,60^ or restoration of short-chain fatty acid availability ^23^.

Several questions remain. Although *EcN* has been administered after antibiotic conditioning to facilitate engraftment ^61–63^, it remains unclear whether NICHE can colonize and remain effective in an intact intestine. Its effects on microbiota recovery and long-term stability also require investigation, particularly because recovering commensals can themselves restore colonization resistance ^26,28,36,37,45^. Finally, defining how *Kp* adapts to sustained metabolic competition may reveal potential escape routes and whether they are constrained by the gut nutrient landscape.

In summary, this study establishes targeted metabolic interference as a rational strategy for controlling intestinal *K. pneumoniae*. By combining transcriptomic inference with metabolomic measurements, we identify conserved *in vivo* nutrient dependencies and distinguish pathway-level fitness effects from their consequences for absolute pathogen expansion. Exploiting these dependencies through engineered competitors and dietary modulation may provide a framework for multilayered interventions against antibiotic-resistant Enterobacteriaceae.

## Limitations of the study

Our study has several limitations. First, all experiments were conducted in female mice, and additional studies in males will be required to determine the generalizability of our findings across sexes. Second, carbon-source consumption by *K. pneumoniae* was assessed in animals subjected to broad-spectrum antibiotic treatment and subsequently colonized with the bacterium. Although anaerobic plating of fecal samples on rich medium did not reveal colonies with morphologies distinct from *K. pneumoniae*, we cannot exclude the presence of low-abundance or non-cultivable microorganisms that may also have contributed to carbon-source consumption. Third, transcriptomic analyses of human samples were compared with *in vitro* reference data generated using MH258 because the corresponding clinical isolates were unavailable for matched experiments. Although the use of a non-isogenic reference would be expected to reduce sensitivity and therefore increase the risk of false negatives rather than false positives, these findings should nevertheless be interpreted with caution. Finally, as this study was intended as a proof of principle, some EcN strains were engineered by plasmid-based expression of *K. pneumoniae* genes. The greater depletion of raffinose and arabitol by NICHE than by EcN-WT provides functional evidence consistent with activity of the introduced metabolic pathways during colonization. However, these observations do not directly establish plasmid retention in EcN-Raf or EcN-Arb over time. Future development of this or similar consortia for translational applications will require direct assessment of genetic stability, and chromosomal integration of the engineered functions may provide a more stable strategy to minimize loss of the introduced traits.

## Resource availability

### Lead contact

Further information and requests for reagents or resources may be directed to and will be fulfilled by lead contact Simone Becattini.

## Materials availability

All materials generated in this study will be made available upon reasonable request.

## Data and code availability

The sequencing data reported in this study are available at the European Nucleotide Archive (ENA) under accession number PRJEB107754. Whole-genome sequencing data generated in this study have been deposited in the NCBI BioProject database under accession number PRJNA1482723. Tabular data have been deposited in Yareta and can be accessed at https://doi.org/10.26037/yareta:mxpjtwxl7rco7nxfpjwz4naxne. The original code is available on GitHub at https://gitlab.unige.ch/becattinilab/zhou_et_al_2026.

## Acknowledgements

We thank all members of our laboratory for useful discussion and critical reading of the manuscript. We thank the iGE3 Genomics Platform of the Faculty of Medicine, University of Geneva, for excellent technical support with sequencing. S.B. is supported by an Eccellenza Professorial Fellowship (PCEFP3_187018) and a Starting Grant (TMSGI3_211235) by the Swiss National Science Foundation. J.K. is supported by funding from Carigest SA, acquired by Dominique Soldati-Favre. Y.Z. is supported by funding from Shanghai Pulmonary Hospital Hundred Talents Program Scholarship. CU and KX are supported by funding from InfectERA-ERANET-through the Acciones complementarias grant (PCIN-2015-094), and by funding from the Spanish MICINN (PID2023-150086OB-I00), and FCT-Portugal grant (InfectERA/0004/2015), respectively.

## Author contributions

Y.Z. and S.B. designed experiments with input from L.C. Y.Z. performed most of the experiments and analyzed data with assistance from C.D. J.P. analyzed and integrated transcriptomic data. T.L. performed analyses. J.K. and O.V.R. performed GC–MS analyses;

I.M. performed LC–MS analyses. M.J.G, C.U and K.B.X. collected and provided human data and provided valuable intellectual input. L.C. performed WGS on optimized EcN and provided critical reagents and discussion. Y.F. supervised and supported Y.Z. S.B. and Y.Z. wrote the paper with input from all authors.

## Declaration of interests

The authors declare no competing interests.

## Experimental model and subject details

### Bacterial strains and growth conditions

*Klebsiella pneumoniae* strains MH258 (carbapenem-resistant, CR-*K. pneumoniae,* CRKp) and ATCC 43816 (hypervirulent, hv-*K. pneumoniae,* hvKp) were used throughout this study. *Escherichia coli* Nissle 1917 (EcN) served as the probiotic chassis strain. All strains used in this study are listed in Table S2.

Bacteria were routinely cultured in Luria–Bertani (LB) medium (BD Difco), or M9 minimal medium (BD Difco) consisting of M9 salts (Na_2_HPO_4_, KHPO_4_, NaCl, and NH_4_Cl), supplemented with MgSO_4_, CaCl_2,_ with glucose or the indicated carbon sources. Glucose was used as the reference carbon source at a final concentration of 0.2% (w/v) in all experiments. Alternative carbon sources were supplemented at the following final concentrations: arabinose −0.2% (w/v), xylose - 0.2% (w/v), glucuronate - 0.2% (w/v), galacturonate - 0.2% (w/v), raffinose - 0.1% or 0.2% (w/v), as indicated, sucrose - 0.1% or 0.2% (w/v), as indicated, gluconate - 0.1% or 0.2% (w/v), as indicated, sorbitol - 0.5% (w/v), arabitol - 0.5% (w/v), oligogalacturonate - 0.5% or 1% (w/v), as indicated, and threonine (30mM). For experiments involving ethanolamine, cultures were supplemented with 40 mM ethanolamine, together with 25 mM vitamin B12, to support ethanolamine catabolism. For experiments involving oligogalacturonate, crude oligogalacturonates were generated by high-temperature (121°C), high-pressure (15 psi) –mediated β-elimination of pectin (pectin from citrus peel, Sigma) and used to supplement the culture medium. All carbon source concentrations used in specific experiments are indicated in following methods section.

For *in vitro* assays modeling dietary inputs, M9 minimal medium was supplemented with 5% (w/v) of the indicated mouse diets, including standard diet (SAFE 150), and high-fat diet (SAFE U8954+1.25% Cholesterol). Diet-conditioned media were prepared using M9 minimal medium as the base. The pH was adjusted to 2.0, and the indicated mouse diets were added and incubated at 37 °C for 30 min to allow complete digestion. After digestion, the pH was readjusted to 7.0, and the media were sterilized by autoclaving prior to use.

For *ex vivo* growth assays, cecal contents were collected from antibiotic-treated mice maintained on the indicated diets for 7 days. Cecal material from three mice per group was pooled, weighed, and resuspended in sterile PBS to generate a uniform cecal-content slurry. The resulting supernatant was used as *ex vivo* cecal-content medium for bacterial growth assays.

Aerobic cultures were shaken at 220 rpm at 37°C. Plates were grown at 37°C for culture streaks and placed at room temperature for colony enumeration. Anaerobic cultures were cultivated in a vinyl anaerobic chamber (Coy Laboratory Products, gas composition: N_2_ 85%, CO_2_ 10%, H_2_ 5%). Antibiotics were provided at the following concentrations: apramycin (100 μg/mL), hygromycin (100 μg/mL), streptomycin (200 μg/mL), rifampin (50 μg/mL), carbenicillin (100 μg/mL), and neomycin (50 μg/mL).

### Mice

WT mice were purchased as JAX C57BL/6J from Charles River, France. All animals were bred and maintained in a specific pathogen-free (SPF) facility at the Medical Center (CMU) of the University of Geneva, Switzerland. Animals were housed in sterile, autoclaved cages with irradiated food (SAFE-150 Diet, or SAFEU8954 supplemented with 1.5% cholesterol, SAFE ®) and autoclaved water available ad libitum. Mouse handling and cage changes were performed by investigators equipped with gloves, masks, shoe covers, and sterile gowns. Unless stated otherwise, all mice used for this study were females of 6–9 weeks of age. All animal experiments were conducted in accordance with legal and ethical regulations. Experimental procedures were approved by the Department of Animal Experimentation of the Geneva canton under license number GE598.

### *K. pneumoniae* reconstitution in mice

All *in vivo* experiments followed the same protocol unless specified otherwise. Naïve gut microbiomes were depleted using an AMNV regimen (ampicillin, metronidazole, neomycin, vancomycin, all at 0.5 g/L in drinking water for 4 days, followed by an additional 2-day vancomycin and metronidazole “flush-out” regimen. After this flush-out period, mice were maintained on vancomycin and metronidazole for the duration of the experiment to ensure sustained depletion of the resident microbiota. Gut microbiome depletion was confirmed by plating fecal pellets on Columbia blood agar plates under anaerobic environment. After confirmation of microbiota depletion, mice were reconstituted with the indicated *K. pneumoniae* strains (MH258 or ATCC 43816) by oral gavage of 200 μL bacterial suspension, containing 10⁵ CFU for MH258 or 10⁴ CFU for ATCC 43816, as indicated. At the indicated time points, mice were sacrificed and cecal contents were collected for downstream analyses, including RNA extraction and metabolomic profiling.

### TRIzol-based RNA extraction from bacterial cultures

For extraction of total RNA from in vitro–cultured bacteria, including *Klebsiella pneumoniae* and *Escherichia coli* Nissle (EcN), a TRIzol-based method was used for all samples intended for RT–qPCR analysis. Bacterial cultures were grown to the desired growth phase and 1 mL of culture was harvested by centrifugation. The bacterial pellet was resuspended in 1 mL of TRIzol reagent (Thermo Fisher Scientific) and samples were processed immediately. For cell lysis, the TRIzol suspension was transferred to screw-cap bead tubes containing silica and zircon beads (Invitrogen). Samples were subjected to mechanical disruption using a bead beater (Microbes 2 mL program; speed 4.30, 1 min total with two 15-s breaks between cycles), ensuring symmetric tube placement. Following lysis, 0.2 mL chloroform per 1 mL TRIzol was added to each sample. Tubes were vigorously shaken for 15 s and incubated at room temperature for 10 min. Samples were then centrifuged at 12,000 × g for 15 min at 4°C, resulting in phase separation into a lower phenol–chloroform phase, an interphase, and an upper aqueous phase. The aqueous phase was carefully transferred to a new RNase-free tube. RNA was precipitated by addition of 0.5 mL isopropanol, followed by gentle inversion to mix and incubation for 10 min at 4°C. Samples were centrifuged at 12,000 × g for 10 min at 4°C, and the resulting RNA pellet was washed twice with 1 mL of 75% ethanol. Each wash was followed by centrifugation at 7,500 × g for 5 min at 4°C. After removal of the supernatant, RNA pellets were air-dried for 5–10 min and resuspended in 50 μL RNase-free water. Purified RNA was aliquoted and stored at −80°C until downstream applications.

### Ǫuantitative reverse transcription PCR (RT-qPCR)

Reverse transcription of RNA extracted from bacteria was performed with ǪuantiTect RT Kit (Ǫiagen) according to the manufacturer’s instructions. cDNA from microbe were diluted 10-fold with RNase-free water before the qPCR. qPCR reactions were set up in MicroAmp Optical 96-well plates (Applied Biosystems) by combining 5 μl of PowerUP SYBR qPCR Master Mix (Applied Biosystems), 3 μl of sample, 1.5 μl of RNase-free water and 0.5 μl of appropriate primer stock solution (10 uM). Plates were sealed with MicroAmp Optical Adhesive Film (Applied Biosystems). Reactions were run on a ǪuantStudio 6 PRO system (Applied Biosystems). Cycling conditions used were in accordance with the master mix manufacturer’s guidelines. qPCR data were normalized to the housekeeping gene, relative gene expression was calculated using the 2^−ΔΔCt method, with the control condition normalized to a value of 1 and all experimental conditions expressed as fold change relative to this control. Values represent the mean of technical replicates for each biological sample. All the primers listed in Table S3.

### RNA extraction from bacterial cultures for RNA sequencing

For the in vitro RNA-seq samples*, K. pneumoniae* cultures grown in LB medium to logarithmic phase (OD₆₀₀ = 0.5) or overnight were collected by centrifugation at 4,000 × g for 5 min at 4°C. Pellets were snap-frozen in liquid nitrogen and stored at −80°C until RNA extraction. Total RNA was extracted using the RNeasy PowerMicrobiome Kit (Ǫiagen) following the manufacturer’s protocol, eluted in nuclease-free water, and stored at −80°C.

### RNA extraction from cecal contents

Cecal contents were collected from mice fed with either a standard diet or a high-fat diet at 7 days after *K. pneumoniae* reconstitution. Mice were sacrificed, and cecal contents were immediately removed from the cecal pouch, snap-frozen in liquid nitrogen, and stored until processing. Microbial RNA was extracted using the RNeasy PowerMicrobiome Kit (Ǫiagen) according to the manufacturer’s instructions, eluted in RNase-free water, aliquoted, and stored at −80°C.

### RNA sequencing and analysis

*K. pneumoniae* RNA sequencing libraries were prepared using the Illumina Stranded Total RNA Prep kit with Ribo-Zero Plus according to the manufacturer’s instructions. Libraries were sequenced on an Illumina NovaSeq 6000 platform using single-end 100 bp reads. Raw sequencing reads were processed using the nf-core/rnaseq pipeline (v3.18.0, doi:10.5281/zenodo.1400710), with parameters adapted for prokaryotic organisms and our experimental design (extra_star_align_args=“--alignIntronMax 20”; extra_trimgalore_args=“--hardtrim5 90”; min_mapped_reads=0). Depending on the K. pneumoniae strain used, reads were mapped to the reference genome ATCC 43816 or MH258. Computations were performed at the University of Geneva using the Baobab high-performance computing service. Resulting gene quantifications were processed using the R programming language. Differential gene expression analysis was performed using DESeq2 (v1.48.2). Principal component analysis (PCA) was conducted to assess global transcriptional differences between experimental conditions and to evaluate clustering of biological replicates. Functional enrichment analyses, including Kyoto Encyclopedia of Genes and Genomes (KEGG) pathway enrichment and gene set enrichment analysis (GSEA), were performed using the fgsea package (v1.34.2). Visualization of transcriptomic data, including heatmaps and volcano plots, was carried out using standard R packages. To compare gene expression between K. pneumoniae strains, a table of orthologous genes was generated by reciprocal BLAST, and a universal PanID identifier was assigned to each gene. In addition, KEGG orthology numbers were assigned by submitting gene sequences to the BlastKOALA service (https://www.kegg.jp/blastkoala), enabling association of genes with KEGG pathways.

## Metabolomics analysis

### Cecal content collection for metabolomic analysis

For metabolomic analyses, the mouse model and antibiotic treatment regimen were identical to those used for *in vivo* transcriptomic profiling. Unless otherwise indicated, mice were maintained on either a standard diet (STD) or high-fat diet (HFD) throughout the experiment. Depending on the experimental design, mice were randomly assigned to the indicated treatment groups, including PBS controls, MH258, ATCC 43816, EcN-WT, or the NICHE consortium. On day 7 after bacterial reconstitution, mice were sacrificed and cecal contents were collected immediately from the cecal pouch. Samples were snap-frozen in liquid nitrogen and stored at −80°C until metabolite extraction and subsequent GC–MS and/or LC–MS analyses.

### GC–MS - Sample Preparation

Cecal samples were gently thawed on ice and 50 mg (+/−0.5 mg) cecal material were transferred to new microfuge tubes. Next, 100 µL chloroform were added to each sample followed by 350 µL of a methanol water mix composed of 18 ml methanol, 2.6 ml water and 400 µL of a 1 mM stock of ^13^C_3_/^15^N β-alanine (Cambridge Isotope Laboratories, CNLM-3946) as internal standard. Note that all solvents used for GC– and LC–MS analyses were of high purity, suitable for mass spectrometry. The samples were thoroughly agitated on a sample mixer at 4 °C for one hour, before centrifuging the samples to sediment insoluble material (20,000 ×g, 4 °C, 10 min). The supernatant was transferred to new microfuge tubes containing 200 µL water and samples mixed again at 4 °C for 15 min before centrifugation to separate the phases (20,000 ×g, 4 °C, 10 min). The upper polar phase was transferred to new microfuge tubes and dried sequentially in mass spectrometry inserts (Machery-Nagel, 702813) using a centrifugal evaporator (Savant Speed Vac Concentrator). The extract was further dried and concentrated through addition of decreasing volumes of methanol. To the dried, concentrated metabolite extracts, 40 µL pyridine (Sigma-Aldrich, 270970) were added, containing methoxyamine hydrochloride (20 mg/mL, Sigma-Aldrich, 226904). Methoxmiation was allowed to proceed over night at room temperature, while rocking the samples gently. The next day, 40 µL of N,O-Bis(trimethylsilyl)trifluoroacetamide with 1% trimethylchlorosilane (BSTFA TMCS, TMS, Supelco, B-023) were added for silylation. The samples were gently rocked at room temperature for one hour. To avoid any residual precipitates to interfere with the analysis, the soluble material was transferred with a clean syringe (Hamilton Company) to new mass spectrometry vials with inserts.

### GC–MS – Analysis and Data Processing

Samples were analyzed using an 8890 GC System (Agilent) equipped with a DB5 capillary column (JCW Scientific, 30 m, 250 μm inner diameter, 0.25-μm film thickness), with a 10-m inert duraguard, connected to a 5977B GC/MSD operated in electron impact (EI, 70 eV) mode linked to a 7693A autosampler (Agilent). The GC–MS settings were as follows: Inlet temperature: 270 °C, MS transfer line temperature: 280 °C, MS source temperature: 230 °C and MS quadrupole temperature: 150 °C. The oven temperature gradient during the sample run was as follows: 80 °C (2 min); 80 °C to 140 °C at 30 °C/min; 140 °C to 250 °C at 5 °C/min; 250 °C to 310 °C at 15 °C/min; 310 °C for 2 min. All samples were analyzed in a randomized sequence following injection of 1 μL in split mode (1:20) with a solvent delay of 7 min in scan mode (*m/z* 70–700). Prior to the sample analysis, 19 authentic standards of sugars of interest were analyzed and their retention time, identifier ions and suitable quantifier ions were recorded (Table S4). Additional 29 metabolites (including amino acids, fatty acids, organic acids and others) were confidently identified using authentic standards and the NIST MS Search 2.4 tool integrated in MassHunter Ǫualitative Analysis Software 12.0 (Agilent) (Table S4). One metabolite, which was not definitively identified, was annotated as unknown disaccharide based on its late elution time and the observation of typical disaccharide ions including *m/z* 204, 217 and 361. These 48 metabolites as well as the internal standard were quantified across all samples using MassHunter Ǫuantitative DA Software 12.1 (Agilent), by integrating the quantifier ion at the defined retention time (RT) with a right and left RT delta of 0.06 min. The data was further processed in Excel Software (Microsoft). The signal for each metabolite was normalized to that of the internal standard (*m/z* 250, corresponding to a prominent fragment of ^13^C_3_/^15^N β-alanine containing two labelled atoms) and expressed as relative abundance with abundance of each metabolite of the PBS control set to 1.

### LC-MS - Sample Preparation

As for the GC-MS analysis, 50 mg (+/-0.5 mg) of cecal sample were transferred to new microfuge tubes. To this, 1 mL extraction solution (4:1 methanol/water, containing 1 µM CHES as technical internal standard for monitoring acquisition) was added to each tube. The samples were thoroughly agitated on a sample mixer at 4 °C for one hour, before centrifuging the samples to sediment insoluble material (20,000 ×g, 4 °C, 10 min). The supernatant was transferred to new microfuge tubes and dried in a centrifugal evaporator (Savant Speed Vac Concentrator). The dried extract was reconstituted in 120 µL of an acetonitrile/water mix (7:3) on a sample mixer at 4 °C for 20 min. To avoid transfer of any insoluble precipitates, the reconstituted samples were centrifuged (20,000 ×g, 4 °C, 10 min). Due to the high concentration of the samples, 10 µL of the supernatants were diluted with 90 µL of an acetonitrile/water mix (7:3, 1:10 *v/v* dilution) and transferred into LC-MS polypropylene vials (Waters, MaxPeak HPS vials, ref 186009186). Ǫuality control (ǪC) samples were constituted by pooling 20 µL of each sample and aliquoted in 2 ǪC samples for monitoring the stability of the acquisition across the sequence, 1 ǪC sample for conditioning the system prior the biological samples and 1 diluted ǪC (dilution 1:1 v/v with reconstitution solvent) for discriminating contaminants.

### LC-MS – Analysis and Data Processing

Samples were measured on a Vanquish Horizon LC system (Thermo Fisher, Waltham, MA, USA) with a zicHilic Premier LC column (2.1 × 100 mm, Waters, Milford, MA, USA) hyphenated to a high-resolution Exploris 120 orbitrap mass spectrometer (Thermo Fisher), following a method adapted from previous studies ^64,65^. The mobile phase A was 20 mM ammonium acetate in water at pH = 9.2 adjusted with ammonium hydroxide, mobile phase B pure acetonitrile and mobile phase C (for column washing only) 200 mM ammonium acetate in water. The elution gradient ran with a flow rate of 0.4 mL/min from 90% to 40% B in 12 min, followed by washing cycles comprising 1 column volume of mobile phase A, 2 volumes of mobile phase C and 3 volumes of mobile phase A. Re-equilibration was performed at the initial conditions for 7.4 column volumes. The LC flow was diverted to waste at 13.5 min. The column oven temperature was set at 30°C and the injection volume was 2 µL. The mass spectrometer was operating in negative mode (−2.3 kV), with internal mass calibration enabled at the start of each run. The source settings were 50 AU for the sheath gas flow, 10 AU and 350°C for the auxiliary gas, 2 AU for the sweep gas, and the ion transfer tube temperature was set at 320°C. S-lens RF was set at 50. Each sample was acquired in centroid mode with an alternating full scan at a 60,000 resolution on a mass range of 80–800 *m*/*z* and top 4 DDA scans at 15,000 resolutions with a normalized collision energy of 15 eV. The AGC target was set at 150%, with automatic accumulation times for MS1 and MS2 scans. MS2 scans were triggered with an intensity threshold of 100,000, an apex window of 45%, and a dynamic exclusion of 3 s. A custom exclusion list based on a blank sample was used to avoid fragmenting contaminants.

Data were pre-processed in MS-DIAL software v5.5.250221 ^66^. Peak detection was performed with 0.01 and 0.025 Da mass tolerances for MS1 and MS2, a minimum peak height of 10,000, a mass slice of 0.05 Da, and smoothing using a linear weighted moving average with smoothing of 3 scans and minimum peak width of 5 scans. Deconvolution of MS2 data was performed with a sigma value of 0.5 and no abundance cutoff. Identification relied on an in-house msp library with a mass tolerance of 0.01 and 0.025 Da (MS1 and MS2) and retention time tolerance of 0.5min. The alignment was performed with a retention time tolerance of 0.05 min and MS1 tolerance of 0.008 Da. Metabolite annotations were filtered for an accurate mass match < 5 ppm, mean intensities in ǪCs > 5 × 10^4^, CVs in ǪCs < 20%, and D-ratios < 40%, following the recommended guidelines ^67^. A total of 78 metabolites (level 1) were retained in the final dataset. We applied PǪN correction based on median ǪC intensities to account for variations in terms of biomass between the samples.

### Construction of isogenic deletion mutants in *K. pneumoniae*

Isogenic gene deletion mutants of *K. pneumoniae* were generated using a CRISPR–Cas9– assisted genome editing approach ^68^. Briefly, single-guide RNAs (sgRNAs) targeting the gene of interest were designed upstream of a 5′-NGG PAM site and cloned into a pSGKP plasmid. Donor DNA templates containing an antibiotic resistance cassette flanked by approximately 500 bp upstream and downstream homology arms were constructed by overlap PCR (Table S3).

First, the plasmid pCasKP, which encodes both Cas9 and the λ-Red recombination system, was introduced into *K. pneumoniae* by electroporation. Overnight cultures of the new bacteria were diluted 1:100 into fresh LB medium and incubated at 30°C with shaking at 220 rpm. Cells were grown for 1 h to an OD600 of ∼0.2, after which expression of the λ Red-recombination systems was induced by addition of L-arabinose to a final concentration of 2.5% (w/v). Cultures were then incubated for an additional 2 h, reaching an OD600 of 0.6–0.8. Cells were subsequently harvested at 4°C, washed extensively with ice-cold sterile water, and resuspended in ice-cold 10% glycerol to generate electrocompetent cells for electroporation. Genome editing plasmids together with donor DNA fragments were introduced by electroporation. Recombinant clones were selected on antibiotic-containing agar plates and verified by PCR. Editing plasmids were subsequently cured by counterselection on sucrose-containing medium and were confirmed by loss of plasmid-associated antibiotic resistance.

Because *K. pneumoniae* encodes two copies of *xylA*, a sequential deletion strategy was employed. One *xylA* locus (peg.558) was first deleted using λ-Red recombination mediated by the pKD46 system. Subsequently, the second *xylA* locus (peg.5240) was deleted using the CRISPR–Cas9 genome editing approach. All procedures followed the same methods described above.

### Construction of complementation plasmids

Complementation plasmids were constructed using the pACYC177 backbone, a low-copy-number plasmid carrying a streptomycin resistance marker, as previously done ^12^. Genes of interest, together with their predicted native promoter regions, were amplified from the *K. pneumoniae* genome and assembled into the plasmid backbone using NEBuilder HiFi DNA Assembly following the manufacturer’s instructions.

Assembled plasmids were transformed into *E. coli* DH5a for propagation and sequence verification, and subsequently introduced into the corresponding *K. pneumoniae* deletion strains by electroporation. Complemented strains were maintained under streptomycin selection.

### Growth curve *in vitro*

For aerobic growth curve assays, *K. pneumoniae* or *EcN* strains were grown overnight in LB medium, The following day, cultures were washed twice with sterile PBS, and cell density was adjusted to an OD600 of 1.0 prior to inoculation into fresh medium. The adjusted bacterial suspension was subsequently diluted 1:100 into fresh M9 minimal medium supplemented with the indicated carbon source for growth curve measurements. Carbon source concentrations were 0.2% for glucose, xylose, gluconate, and raffinose, and 0.5% for arabitol and oligogalacturonate. *In vitro* growth of *Kp* was estimated by measuring every 15 min for 24 h the optical density at 600 nm on a TECAN Infinite 200 PRO (TECAN) in aerobic conditions.

For anaerobic growth curve assays, bacterial cultures were prepared as described for aerobic conditions, with all handling and incubation performed in an anaerobic chamber. OD600 values were measured manually at 1–2 h intervals for 10–12 h, followed by a final measurement at 24 h.

### *In vitro* competition assays

All *in vitro* competition assays were performed under anaerobic conditions. *K. pneumoniae* strains were prepared as described for growth curve assays. Briefly, wild-type strains were mixed at a 1:1 ratio with the corresponding isogeneic deletion mutants or complemented strains, and the mixtures were subsequently diluted 1:100 into M9 minimal medium supplemented with the indicated carbon sources. Carbon source concentrations were: glucose (0.2%, w/v), raffinose (0.1% or 0.2%, w/v), sucrose (0.1% or 0.2%, w/v), and oligogalacturonate (0.5%, w/v). For competition assays performed in mixed-carbon medium, M9 minimal medium was supplemented with raffinose, arabinose, arabitol, xylose, and gluconate, each at a final concentration of 0.1% (w/v). At the specified time points, bacterial abundance was quantified by CFU enumeration on selective media. Competitive indices values were determined by comparing the ratio of mutant (or complemented strain) to wild-type bacteria recovered at each time point, normalized to the corresponding input ratio. Wild-type bacteria served as the reference strain for all competition analyses

### *In vivo* competition assays

The mouse model was established as described for *K. pneumoniae* reconstitution. In experiments involving MH258, mice were continuously maintained on ampicillin-containing drinking water following microbiota depletion and throughout *K. pneumoniae* colonization.

Wild-type, mutant, and the corresponding complemented mutant *K. pneumoniae* strains were grown to early exponential phase, washed, and resuspended in cold PBS. Equal numbers of wild-type and mutant bacteria were mixed at a 1:1 ratio (10^5^ CFU in total), and mice were orally gavaged with 200 μL of the bacterial suspension. The input ratio was confirmed by plating serial dilutions of the inoculum.

Fecal samples were collected at the indicated time points post-gavage, serially diluted, and plated on selective media to enumerate wild-type and mutant colonies. Competitive indices were calculated as the ratio of mutant to wild-type CFU recovered from fecal samples, normalized to the input ratio.

### Human fecal sample collection and analysis

Fecal samples were collected from three hospitalized patients receiving antibiotic treatment at the Hospital La Fe (Valencia, Spain) (see Table S1 for details). All patients were under treatment for acute myeloid leukemia and colonized with *K. pneumoniae* at the time of sampling. The protocol for sample collection was approved by the Ethics Committee of CEIC Dirección General de Salud Pública y Centro Superior de Investigación en Salud Pública (20130515/08). All patients gave their consent for the collection and use of samples. Participants did not receive any compensation for participating in the study.

Patient 1 was treated with meropenem and teicoplanin. Two fecal samples were collected 16 days apart. The first fecal samples was collected 9 days after antibiotic treatment initiation and yielded to the isolate CU402. The second fecal sample was obtained 25 days after meropenem initiation and 9 days after teicoplanin cessation. This sample yielded to CU406 isolate. Patient 2 received ciprofloxacin, and a fecal sample was collected 24 days after ciprofloxacin initiation. This sample yielded to the *K. pneumoniae* CU459 isolate. Patient 3 was treated with meropenem, and a fecal sample was collected 10 days after treatment initiation yielding to *K. pneumoniae* strain CU462. Samples were resuspended in RNAlater before freezing them at −80°C.

In addition, all four *K. pneumoniae* isolates (CU402, CU406, CU459, and CU462) were subjected to Illumina-based whole-genome sequencing, and the resulting assemblies were used as strain-specific reference genomes for transcriptomic read alignment and downstream gene expression analyses.

Microbial RNA was extracted directly from fecal samples using the same procedures described for mouse cecal contents and subjected to RNA sequencing to profile *K. pneumoniae* gene expression in the human intestinal environment. Transcriptomic data processing and differential expression analyses were performed using the same pipeline as described for the mouse colonization experiments.

To assess overall microbial community composition, total DNA was also extracted from fecal samples for 16S rRNA gene sequencing. Operational taxonomic unit (OTU)–based analyses were performed to evaluate bacterial community diversity and to estimate the relative abundance of *K. pneumoniae* within each sample, as previously described ^69^.

## Optimization of *Escherichia coli* Nissle 1G17 for enhanced utilization of specific carbon sources

### Promoter and RBS replacement–based overexpression of carbon utilization genes

As an initial strategy, EcN was engineered to constitutively overexpress key transporters and metabolic enzymes involved in gluconate and arabinose utilization. Specifically, genes encoding the gluconate transporter (*gntT*) and metabolic enzyme (*gntK*), as well as the arabinose transporter (*araF*) and metabolic enzyme (*araB*), were targeted. Genome editing was performed using the same CRISPR–Cas9–based strategy described for *K. pneumoniae* gene deletion, with the modification that the native promoters were replaced by a strong synthetic promoter (J23199) and a strong ribosome-binding site (B0034) (https://parts.igem.org/Main_Page) supplied within the donor DNA sequence. This approach enabled constitutive high-level expression of the targeted genes. All primers used for genetic manipulation are listed in Table S3.

### *In vivo* experimental evolution of EcN for improved carbon source utilization

To enable *in vivo* selection of EcN variants with enhanced carbon utilization, a competitive experimental evolution strategy was employed. Using the same genome editing approach described above, two EcN strains were generated by inserting distinct antibiotic resistance markers (apramycin or hygromycin resistance) into the intergenic region between *rhtB* and *rhtC*, a chromosomal locus previously described as a neutral insertion site ^70^ in *E. coli*.

Antibiotic-treated mice were orally gavaged with an equal mixture (1:1) of the two marker-tagged EcN strains and supplied with drinking water containing 0.6% (w/v) of the indicated carbon source. Fecal samples were collected longitudinally, and changes in the relative abundance of the two EcN populations were monitored by selective plating. Representative colonies from the enriched population were subsequently isolated and evaluated by *in vitro* competition assays against the parental EcN strain under anaerobic conditions in M9 minimal medium containing the corresponding carbon source as the sole carbon source. Clones that consistently exhibited a competitive advantage over the parental strain were selected as the final carbon source–optimized EcN strains for subsequent characterization and *in vivo* studies. Clones showing consistent enrichment were isolated and designated as carbon source–optimized EcN strains. Primers used for strain construction are listed in Table S3.

To define the genetic basis of the advantageous clones obtained during *in vivo* evolution, the EcN-OGA and EcN-Ara strains were subjected to whole-genome sequencing. Whole-genome sequencing of the bacterial isolates was performed using the Illumina NovaSeq 6000 platform. Genomic DNA libraries were prepared according to standard Illumina protocols and sequenced to generate 150-bp paired-end reads. The raw reads were quality filtered using Trimmomatic v0.39 ^71^, followed by de novo assembly using SPAdes v3.15 ^72^ with default settings to generate draft genomes. In addition, the trimmed reads were mapped to the corresponding Nissle 1917 reference genome (NZ_CP058217) using BWA v0.7.19 with default parameters ^73^. Single nucleotide variants (SNVs) were identified using FreeBayes v1.3.10 (https://github.com/freebayes/freebayes) and annotated using SnpEff v5.4c ^74^.

### Heterologous expression of *K. pneumoniae* raffinose and arabitol operons in EcN

To enhance EcN utilization of raffinose and arabitol, plasmids carrying the corresponding raffinose operon (*rafB*-*cscA*-*rafD*) or arabitol operon (*dalT*-*dalD*-*atlB*) derived from *K. pneumoniae* were introduced into EcN. Genes were cloned into the low-copy-number plasmid pACYC177, and plasmid construction followed the same strategy used for complementation plasmids described above. The resulting plasmids were transformed into EcN WT, generating EcN strains optimized for raffinose or arabitol utilization. All cloning primers are listed in Table S3.

### Deletion of the *gntR* repressor to enhance gluconate utilization

As an additional strategy to improve gluconate utilization, an isogenic *gntR* gene deletion mutant was constructed in EcN. The *gntR* transcriptional repressor was deleted using the same CRISPR–Cas9–based genome editing approach employed for *K. pneumoniae* deletion mutants. Removal of *gntR* resulted in de-repression of gluconate utilization genes, thereby enhancing EcN growth on gluconate as the sole carbon source. All primers used for genome editing are provided in Table S3.

### Generation of barcoded EcN strains

To enable longitudinal tracking of the relative abundance of individual EcN strains within the NICHE consortium during intestinal colonization, unique 30-bp DNA barcodes were introduced into the optimized EcN strains. Barcode insertion was performed for EcN-Ara, EcN-Raf, EcN-OGA, EcN-Xyl, and EcN-GlcA, whereas EcN-Arb remained unmodified and could be distinguished from the other strains during downstream analyses.

Each barcode, together with an ampicillin resistance cassette, was inserted into the neutral chromosomal locus located between the *rsmI* and *agaI* genes using the CRISPR-Cas9-mediated homologous recombination strategy described above. Correct integration of the barcode constructs was confirmed by PCR. Primer sequences used for strain construction and verification are listed in Table S3.

### Functional validation of optimized EcN strains

All experimental procedures were performed using the same conditions and workflows as those described above for *K. pneumoniae*, unless otherwise specified. The functional consequences of EcN optimization were assessed using a combination of growth curve analyses, *in vitro* competition assays, and RT-qPCR, as indicated in the corresponding figure legends.

For growth assays, optimized EcN strains and EcN WT were cultured in M9 minimal medium supplemented with glucose or the relevant carbon source under aerobic or anaerobic conditions, and growth was monitored by OD600 measurements.

To directly compare competitive fitness, *in vitro* competition assays were performed under anaerobic conditions by co-culturing optimized EcN strains with EcN WT at a 1:1 ratio in M9 medium containing the indicated carbon source. Carbon source concentrations were 0.2% for glucose and arabinose, 0.1% for gluconate, raffinose, and xylose, 0.5% for arabitol, and 1% for oligogalacturonate. Bacterial abundance was quantified by CFU enumeration at defined time points, and competitive indices were calculated relative to EcN WT, as described above.

In addition, RT-qPCR was performed to assess transcriptional changes in carbon utilization genes. Bacterial cultures were harvested at mid-logarithmic phase under the indicated growth conditions, RNA was extracted, and gene expression levels were quantified relative to EcN WT. Carbon source concentrations was the same as growth curve.

### *In vitro* protection and decolonization assays with optimized Nissle and *K. pneumoniae* MH258

All *in vitro* competition assays were performed in minimal M9 medium supplemented with glucose or individual carbon sources and incubated under anaerobic conditions. Glucose (0.2%, w/v) was included as the reference carbon source. For assays performed with alternative carbon sources, concentrations were: arabinose (0.1%, w/v), xylose (0.1%, w/v), gluconate (0.1%, w/v), raffinose (0.1%, w/v), arabitol (0.5%, w/v), and oligogalacturonate (0.5%, w/v).

### Protection model (pre-inoculation of optimized Nissle)

To assess whether optimized Nissle strains could prevent *K. pneumoniae* expansion, M9 medium containing glucose or the indicated carbon source was inoculated with PBS, EcN WT, or the corresponding optimized Nissle at 10⁷ CFU. Cultures were incubated for 10 h, after which *K. pneumoniae* MH258 was introduced at 10⁵ CFU. Bacterial burdens were quantified at the indicated time points by selective plating to enumerate EcN and MH258.

### Decolonization model (post-establishment introduction of optimized Nissle)

To assess whether optimized Nissle strains could suppress MH258 following prior establishment, M9 medium containing glucose or the indicated carbon source was first inoculated with *K. pneumoniae* MH258 at 10⁵ CFU and allowed to establish. Subsequently, optimized Nissle, EcN WT (each at 10⁹ CFU), or PBS was added. Bacterial burdens were quantified at the indicated time points by selective plating.

## Assembly of the NICHE consortium and *in vivo* protection and therapeutic experiments

### Assembly of the NICHE consortium

To generate a defined probiotic consortium optimized for competitive utilization of gut carbon sources, six EcN strains individually optimized for utilization of arabinose, arabitol, raffinose, oligogalacturonate, gluconate, and xylose were assembled into a single consortium, termed NICHE (Nissle Integrated Consortium for Heterogeneous Energy utilization). Each optimized EcN strain was grown separately under standard conditions, and equal CFUs of each strain were combined immediately prior to administration to mice. The final NICHE inoculum thus contained an equal representation of all six optimized EcN strains.

### *In vivo* protection model

For protection experiments, the mouse model and antibiotic treatment regimen were identical to those described above for *K. pneumoniae* reconstitution. Briefly, antibiotic-treated mice maintained on either a standard diet (STD) or high-fat diet (HFD), as indicated, were orally gavaged with a single dose of NICHE, EcN-WT, or an equal volume of PBS as a control. Following reconstitution, mice were allowed to establish stable EcN colonization.

Four days after EcN or NICHE administration, mice were orally challenged with *K. pneumoniae* (MH258, 10^5^ CFU; or ATCC 43816, 10^4^ CFU, as indicated). Fecal samples were collected longitudinally at the indicated time points, and *K. pneumoniae* and EcN burdens were quantified by selective plating.

For experiments using ATCC 43816, mice were euthanized on day 3 after pathogen challenge. Cecal tissue, mesenteric lymph nodes (MLNs), liver, and spleen were aseptically collected for bacterial enumeration. To distinguish tissue-associated bacteria from bacteria adherent to the cecal surface, each cecum was divided into two equal portions. One portion was homogenized directly, whereas the other was incubated in PBS containing 100 μg/mL gentamicin for 1 h at 37°C, followed by extensive washing with sterile PBS before tissue homogenization and CFU enumeration.

### *In vivo* therapeutic model

For therapeutic experiments, antibiotic-treated mice were first orally gavaged with *K. pneumoniae* to establish intestinal colonization. After 3 days, when *K. pneumoniae* burdens had reached a stable peak, mice were treated by oral gavage with NICHE, EcN WT, or PBS.

Following treatment, fecal samples were collected at the indicated time points, and *K. pneumoniae* burdens were quantified by selective plating. This model was used to evaluate the capacity of NICHE to reduce or eliminate established *K. pneumoniae* colonization in the intestinal tract.

### Ǫuantification of barcoded EcN strains by qPCR

To determine the relative abundance of individual EcN strains within the NICHE consortium during intestinal colonization, mice were orally gavaged with the barcoded NICHE consortium. Fecal samples were collected on days 1, 7, and 14 after inoculation, and total genomic DNA was extracted using the PureLink™ Microbiome DNA Purification Kit (Invitrogen) according to the manufacturer’s instructions.

For each barcoded strain, a standard curve was generated by correlating DNA concentration with the corresponding Ct value using barcode-specific primer pairs. Purified fecal DNA samples were then subjected to quantitative PCR (qPCR) using the same primer sets. The abundance of each strain was calculated based on its corresponding standard curve and expressed as the relative abundance within the total NICHE population. Primer sequences are listed in Table S3.

### Statistical analysis

Statistical analyses were performed using GraphPad Prism and R, as specified in the corresponding figure legends. RNA-seq differential expression analyses were conducted using DESeq2, with principal component analysis (PCA) and KEGG pathway enrichment performed using standard R packages. Growth curve, RT-qPCR, and metabolomic data are presented as mean ± SD and were analyzed using unpaired two-tailed t tests for comparisons between two groups. Competitive indices from in vitro and in vivo competition assays are presented as geometric means and were analyzed using one-sample *t* tests relative to a value of 1. For *in vivo* competition assays, competitive-index values were log10-transformed prior to statistical analysis. Comparisons against the neutral value (competitive index = 1) were performed using two-sided one-sample *t* tests, whereas direct comparisons between mutant and complemented mutant strains or comparions between standard diet and high-fat diet groups were performed using two-sided unpaired Welch’s t-tests. CFU data were log10-transformed prior to statistical analysis because bacterial counts typically spanned several orders of magnitude; log transformation was used to reduce skewness and improve variance stabilization for parametric statistical comparisons. Mouse colonization data are shown as geometric means ± geometric SD, and comparisons between two groups were performed using unpaired two-tailed *t* tests on log10-transformed CFU values. For experiments involving more than two groups or multiple time points, statistical significance was determined using one-way ANOVA followed by Tukey’s multiple comparisons test. Statistical significance thresholds are indicated in the figure legends on log10-transformed CFU values where applicable.

## Notes

### Competing Interest Statement

The authors have declared no competing interest.

