## Supplementary Information for "Defining nutrient dependencies of intestinal *Klebsiella pneumoniae* guides rational probiotic engineering"

**Supplemental information**

### Transcriptome-guided rational probiotic engineering exploits metabolic interference against intestinal *Klebsiella pneumoniae*

Ying Zhou, Clement David, Grishma Kulkarni, Tengfei Long, María J. Garzón, Julien Prados, Joachim Kloehn, Olivier Von Rohr, Isabel Meister, Karina B. Xavier, Carles Ubeda, Fangyou Yu, Liang Chen, Simone Becattini

Includes Supplementary Figures 1-14

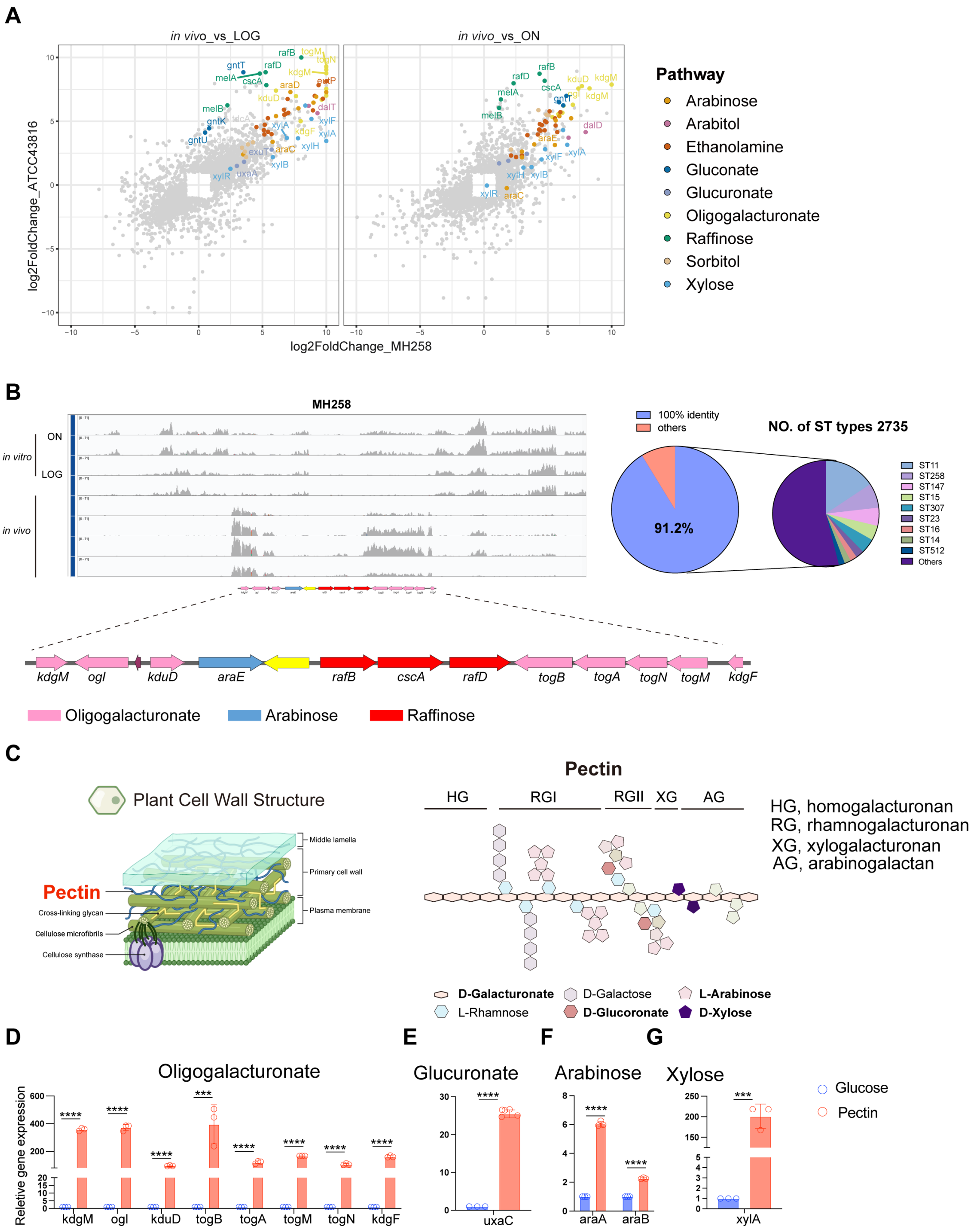


**Figure S1. Conserved carbon metabolism programs in *K. pneumoniae* during intestinal colonization. (A)** Correlation analysis of differential gene expression between MH258 and ATCC 43816. Relative expression values are shown as *in vivo* relative to *in vitro* comparisons. Each dot represents one gene. Colors indicate different carbohydrate metabolic pathways. *n* = 3 or 4 biological replicates per group. **(B)** Visualization of transcriptomic read coverage of MH258 under *in vitro*

(top) and *in vivo* (bottom) conditions using Integrative Genomics Viewer (IGV) (left). The highlighted region represents a 13-gene island that is significantly upregulated *in vivo*. Genes within this island are primarily associated with oligogalacturonate (pink), arabinose (blue), and raffinose (red) metabolism. To assess conservation of this gene island, 76,609 *K. pneumoniae* genomes were retrieved from the NCBI database (right). The pie chart indicates that 91.2% of genomes carry this gene island with 100% sequence identity (purple). These strains are distributed across 2,735 distinct sequence types (STs), with different colored segments representing different ST groups. **(C)** Schematic illustration of plant cell wall composition (left), highlighting pectin as a major structural component, and the molecular composition of pectin (right). Carbon sources identified as utilized by *K. pneumoniae in vivo* based on transcriptomic analysis, including oligogalacturonate, glucuronate, arabinose, and xylose, are major constituents of pectin. **(D-G)** RT–qPCR analysis of *K. pneumoniae* gene expression in media supplemented with heat-depolymerized pectin, to release individual components. Data are shown as mean ± SD from three independent experiments. Statistical significance was determined using unpaired two-tailed *t* tests (*$P < 0.05$; **$P < 0.01$; ***$P < 0.001$; ****$P < 0.0001$).

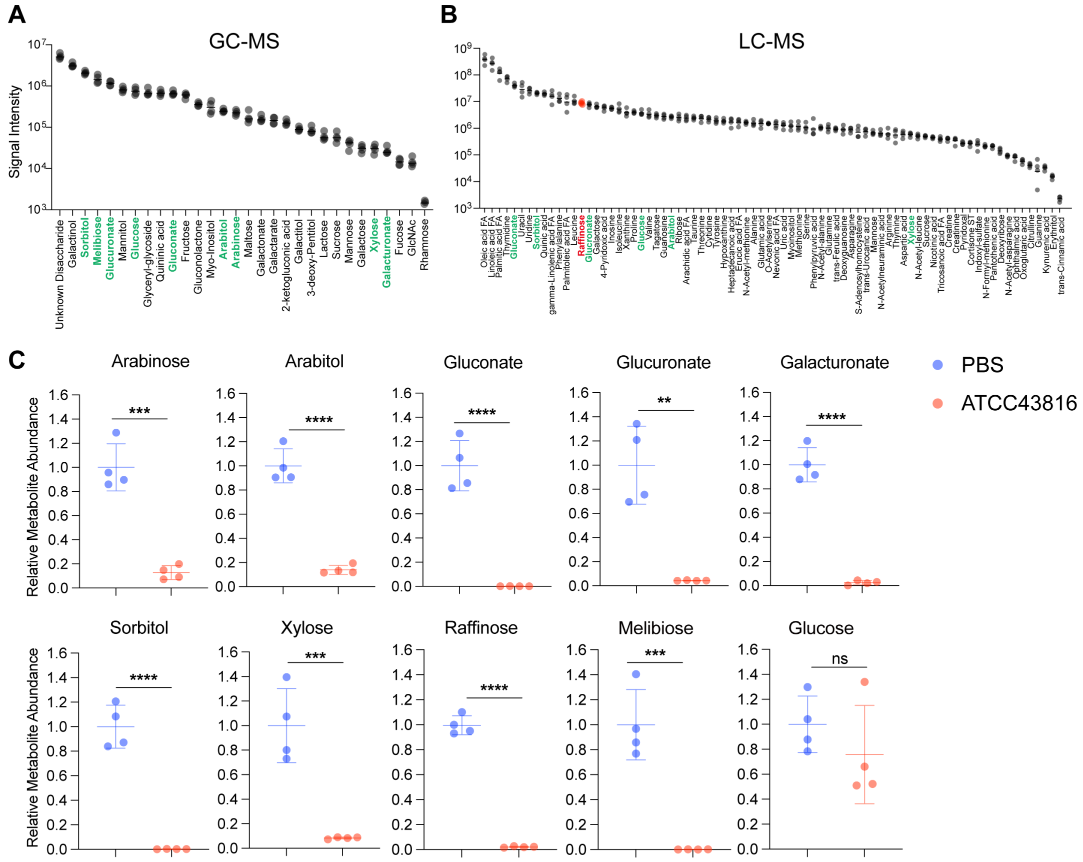


**Figure S2. Metabolomic profiling of cecal contents identifies carbon sources utilized by hypervirulent *K. pneumoniae*. (A-B)** Signal intensities of selected metabolites in cecal contents from antibiotic-treated mice, measured by GC– (A) and LC–MS (B). Carbon sources highlighted in green correspond to those predicted to be utilized by *K. pneumoniae in vivo* based on transcriptomic data. Raffinose (red) is a trisaccharide that cannot be detected by GC–MS and was therefore quantified using LC–MS. Each dot represents one mouse, and data are shown as geometric mean with geometric SD. **(C)** GC–MS and LC-MS–based quantification of individual metabolites in the cecal content from ABX (antibiotics)-treated mice one week after reconstitution with *K. pneumoniae* ATCC43816, compared to PBS-treated controls. Each dot represents one mouse (n=3); horizontal lines indicate means ± SD. Note: oligogalacturonate and raffinose cannot be directly detected by GC–MS; melibiose is here used as a proxy for raffinose abundance, as it is generated upon raffinose degradation either enzymatically or during GC–MS analysis. Raffinose abundance was quantified directly by LC–MS. Statistical significance was determined using unpaired two-tailed *t* tests (ns, Non-significant, *$P < 0.05$; **$P < 0.01$; ***$P < 0.001$; ****$P <$ 0.0001).

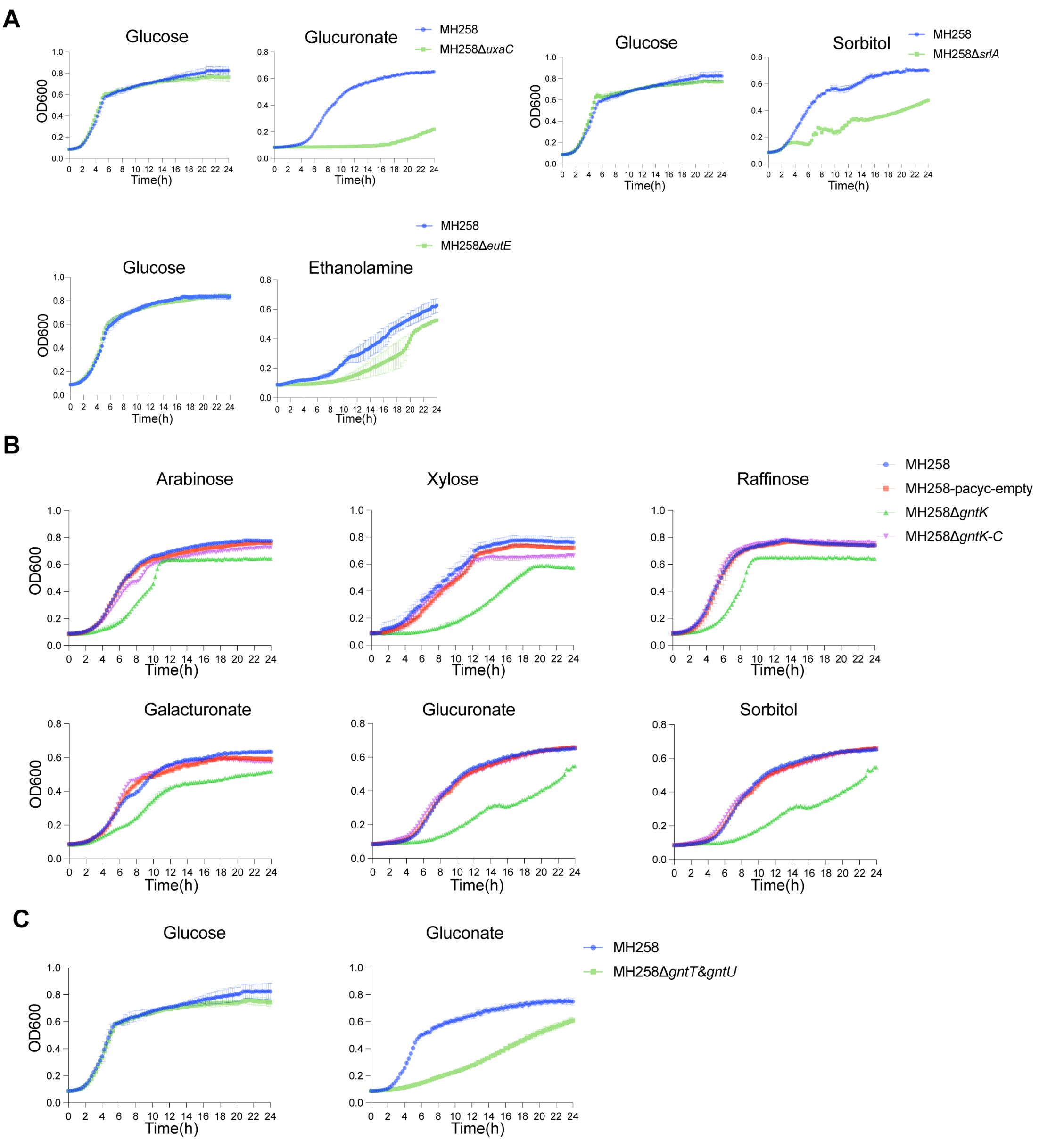


**Figure S3. Growth defects of gluconate and sugar metabolism mutants in *K. pneumoniae*. (A)** Aerobic growth curves of wild-type and mutant strains in M9 minimal medium supplemented with glucose or the indicated carbon sources. Carbon source concentrations were 0.2% for glucuronate, 0.5% for sorbitol, and 40 mM for ethanolamine (with 25 mM vitamin B12 supplementation). Data are shown as mean ± SEM from three independent experiments. **(B)** Growth curves of the MH258 wild-type strain, the *gntK* deletion mutant, and the complemented strain in M9 minimal medium supplemented with the indicated carbon sources. Carbon source concentrations were 0.2% for arabinose, xylose, galacturonate, glucuronate, and raffinose, and 0.5% for sorbitol. Data are shown as mean ± SEM from three independent experiments. **(C)** Growth curves of the MH258 wild-type strain and the *gntT/U* double-deletion mutant in M9 minimal medium supplemented with gluconate (0.2%). Data are shown as mean ± SEM from three independent experiments.

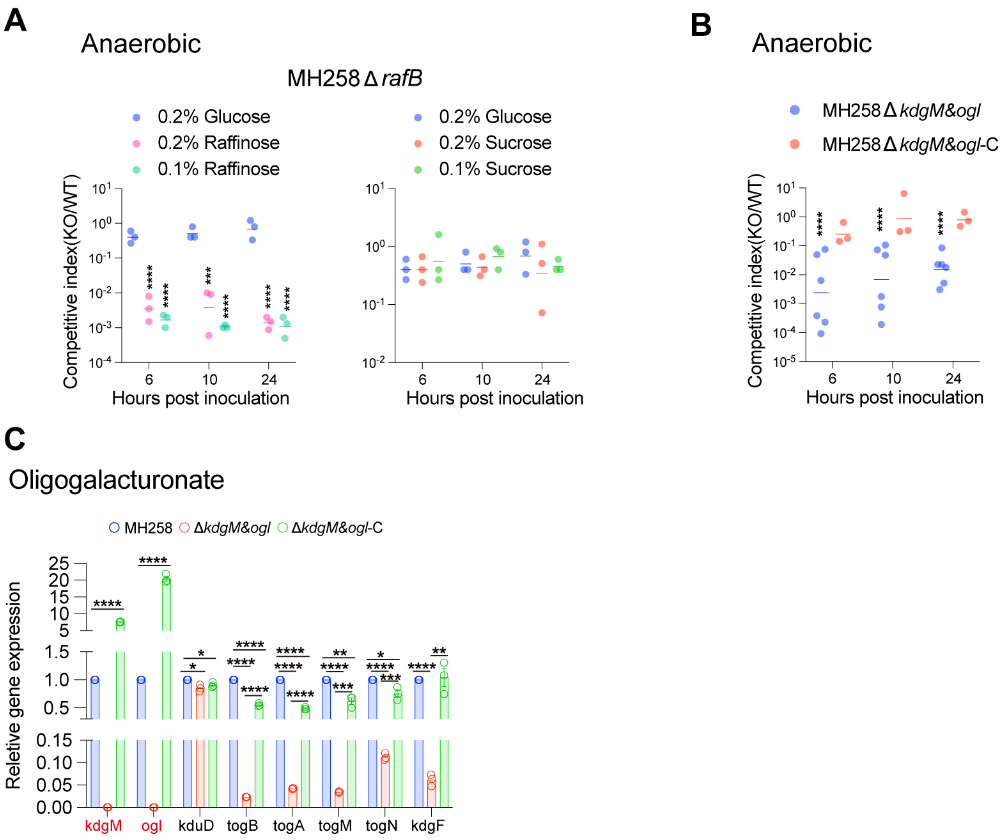


**Figure S4. Functional validation of *K. pneumoniae* mutants involved in raffinose and oligogalacturonate utilization. (A)** Competition assays under anaerobic conditions between the MH258 wild-type strain and *rafB* deletion mutant in M9 minimal medium supplemented with glucose, raffinose, or sucrose at the indicated concentrations. Competitive indexes were determined based on selective plating at the indicated time points. Results are shown as geometric means from three independent experiments. Statistical significance was assessed using one-sample *t* tests (**P* < 0.05; ***P* < 0.01; ****P* < 0.001; *****P* < 0.0001). **(B)** Competition assay under anaerobic conditions assessing oligogalacturonate utilization. Competitions were performed between the MH258 wild-type strain, the *kdgM/ogl* DKO deletion mutant, and the complemented strain in M9 minimal medium supplemented with oligogalacturonate (0.5%). Competitive indexes were determined based on selective plating at the indicated time points. Results are shown as geometric means from three or six independent experiments. Statistical significance was assessed using one-sample *t* tests (**P* < 0.05; ***P* < 0.01; ****P* < 0.001; *****P* < 0.0001). **(C)** RT–qPCR analysis of genes associated with oligogalacturonate metabolism in the MH258 wild-type strain, the *kdgM/ogl* DKO deletion mutant, and the complemented strain. Strains were grown to logarithmic phase in M9 minimal medium supplemented with oligogalacturonate (0.5%), followed by RNA extraction. Data are presented as mean ± SD from three independent experiments. Statistical significance was determined using unpaired two-tailed *t* tests (**P* < 0.05; ***P* < 0.01; ****P* < 0.001; *****P* < 0.0001).

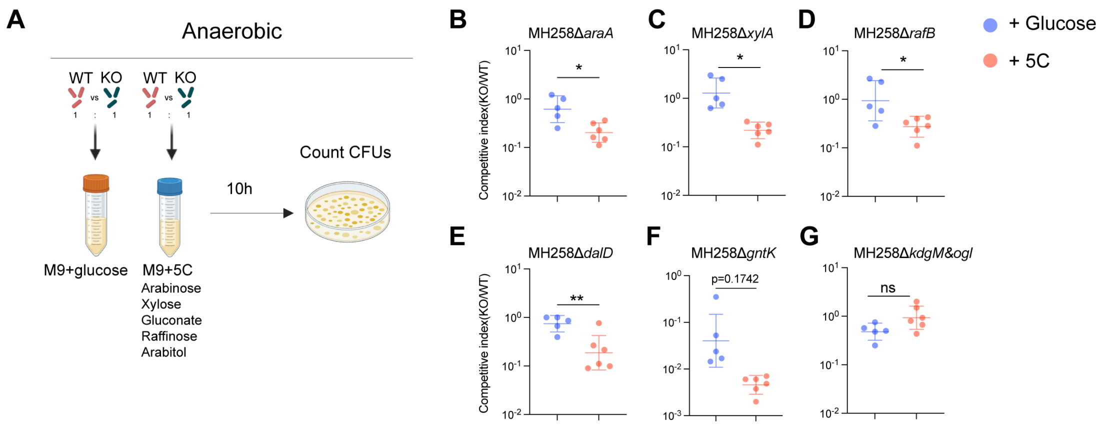


**Figure S5. Individual carbon utilization pathways remain important for *K. pneumoniae* fitness in a mixed-carbon environment. (A)** Schematic of the in vitro competition assay. Wild-type (WT) and the corresponding carbon utilization mutant (KO) were mixed at a 1:1 ratio and cultured anaerobically in M9 minimal medium supplemented with either 0.2% (w/v) glucose (control) or a mixed-carbon medium containing raffinose, arabinose, arabitol, xylose, and gluconate, each at a sub-saturating concentration of 0.1% (w/v). Competitive indices were determined after incubation by comparing the ratio of mutant to WT bacteria. **(B–G)** Competitive indices of the indicated carbon utilization mutants relative to WT in the mixed-carbon medium. Data are presented as geometric means ± geometric SD. Each point represents one biological replicate, and data are pooled from two independent experiments. Statistical significance was determined using an unpaired two-tailed *t*-test (ns, Non-significant, **P* < 0.05; ***P* < 0.01; ****P* < 0.001; *****P* < 0.0001).

**A**

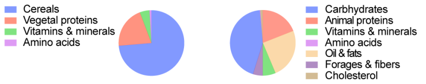


| Macronutrients | SAFE 150 (STD) | SAFE U8954 + 1.25% Cholesterol(HFD) |
|---|---|---|
| Total protein (%) | 18.0 | 17.12 |
| Plant-derived protein (%) | 18.0 | 0 |
| Animal-derived protein (%) | 0 | 17.12 |
| Total fat (%) | 4.8 | 25.41 |
| Total carbohydrates (NFE, %) | 56.9 | 48.76 |
| of which sugars (%) | 3.4 | 34.88 |
| of which starch (%) | 41.0 | 10.25 |
| Crude fiber (%) | 4.1 | 3.5 |
| Ash (%) | 4.2 | 3.2 |
| Moisture (%) | 12.0 | 3.2 |
| Energy density | | |
| Energy density (kcal/g, Atwater) | 3.43 | 4.81 |
| Energy density (kcal/g, ME/Pig) | 3.26 | 4.67 |

**B**

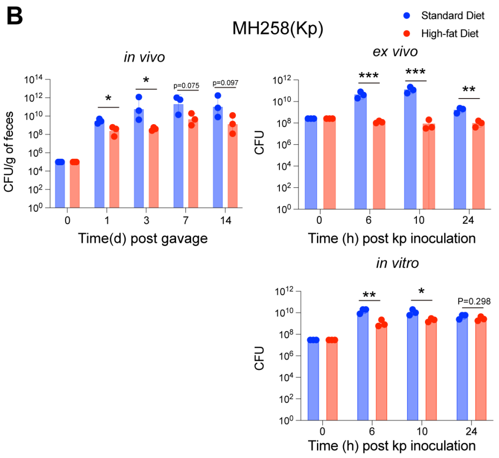


**C**

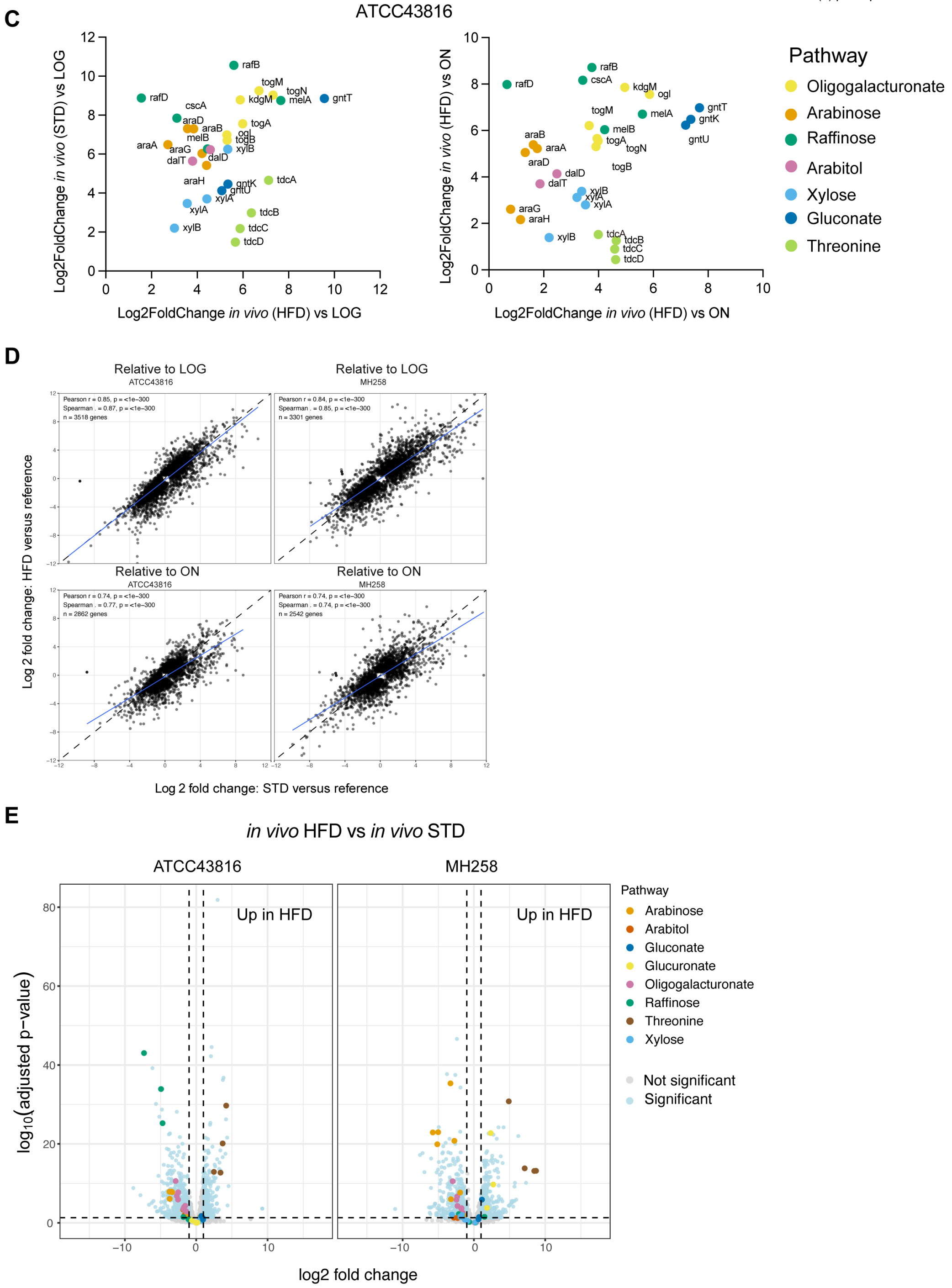

**Figure S6. Dietary composition and its impact on *K. pneumoniae* growth.**

**(A)** Composition of the standard diet (STD) and high-fat diet (HFD) used in this study. Pie charts illustrate the relative contribution of major dietary ingredients, and the accompanying table summarizes macronutrient composition and total dietary energy content of each diet. **(B)** Growth of *K. pneumoniae* MH258 using the two different feeds *in vivo*, *ex vivo*, and *in vitro*. For *in vivo* experiments, bacterial burdens were quantified in mice fed STD or HFD; each dot represents one mouse, and data are presented as geometric mean ± geometric SD. For *ex vivo* assays, cecal contents were collected from antibiotic-treated mice fed STD or HFD for 7 days, pooled from three mice per diet, resuspended, and used as growth media; results are shown as geometric means ± geometric SD from one of three independent experiments. For *in vitro* assays, MH258 was grown in M9 minimal medium supplemented with 5% (w/v) of the indicated mouse diets, dissolved with pH and autoclaved (see Methods); data are presented as geometric mean ± geometric SD from three independent experiments. Statistical significance was determined using unpaired two-tailed *t* tests (*$P < 0.05$; **$P < 0.01$; ***$P < 0.001$; ****$P < 0.0001$). **(C)** Correlation analysis of differential expression of genes associated with seven key carbon sources in *K. pneumoniae* ATCC43816 colonizing mice fed STD versus HFD. Each dot represents one gene, with colors indicating different carbon utilization pathways. **(D)** Pairwise correlation of genome-wide transcriptional responses between mice fed the standard diet (STD) and high-fat diet (HFD). Differential gene expression was calculated relative to logarithmic-phase (LOG) cultures (top) or overnight (ON) cultures (bottom) for ATCC43816 (left) and MH258 (right). Each dot represents one gene. Pearson correlation coefficients (r), corresponding $P$ values, and the number of genes analyzed (n) are shown in each panel. The solid blue line represents the linear regression fit, and the dashed line indicates the line of identity. **(E)** Volcano plots showing differential gene expression of *K. pneumoniae in vivo* in mice fed a high-fat diet (HFD) relative to a standard diet (STD) for ATCC43816 (left) and MH258 (right). Light blue dots indicate significantly differentially expressed genes ($|\log_2$

fold change| > 1, adjusted $P < 0.05$), whereas gray dots indicate genes that did not meet the significance threshold. Genes involved in the indicated carbon utilization pathways (arabinose, arabitol, gluconate, glucuronate, oligogalacturonate, raffinose, threonine, and xylose metabolism) are highlighted in different colors as shown in the legend.

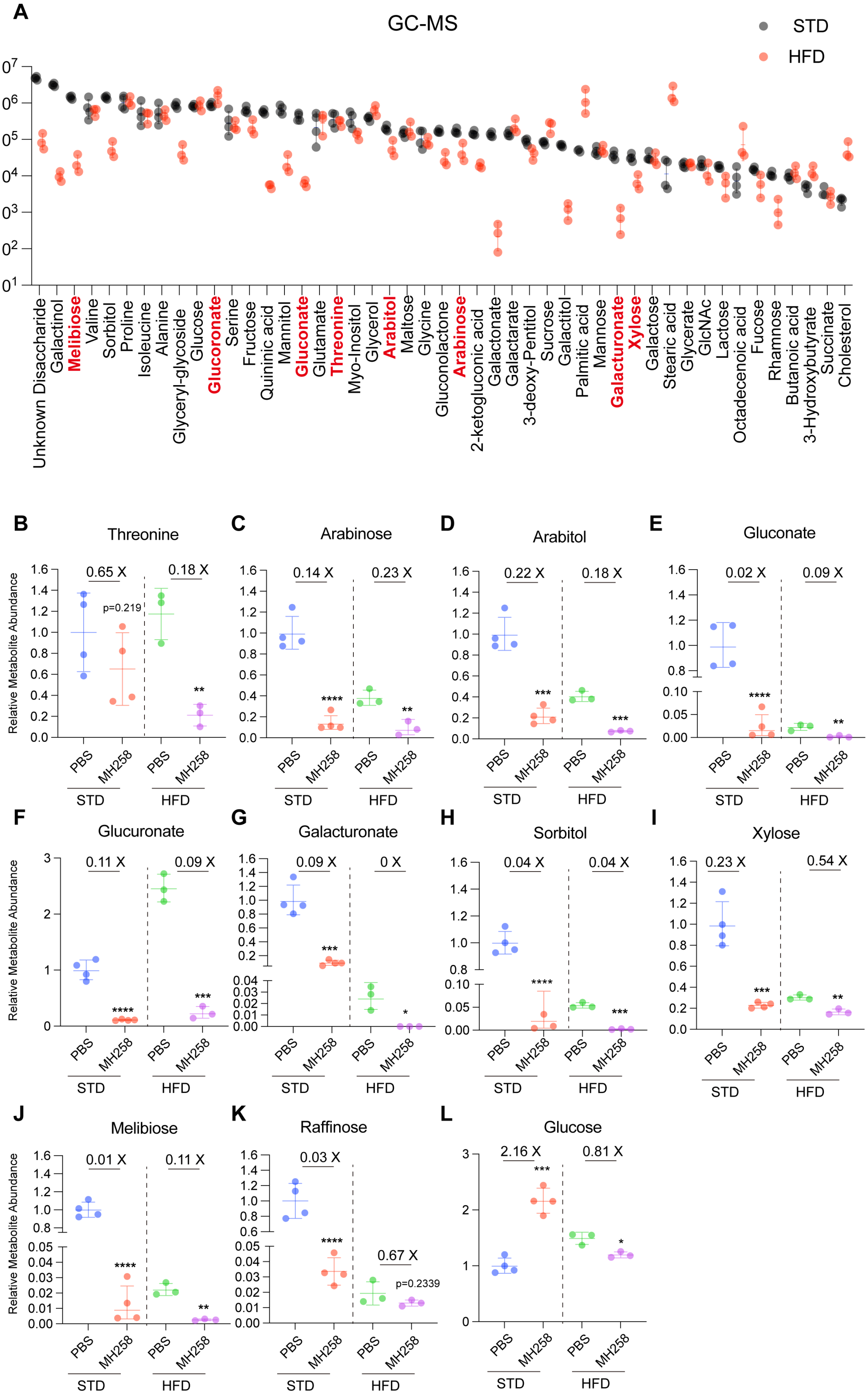
A
GC-MS
STD
HFD
Unknown Disaccharide
Galactinol
Melibiose
Valine
Sorbitol
Proline
Isoleucine
Alanine
Glyceryl-glycoside
Glucose
Glucoronate
Serine
Fructose
Quininic acid
Mannitol
Gluconate
Glutamate
Threonine
Myo-Inositol
Glycerol
Arabitol
Maltose
Glycine
Gluconolactone
Arabinose
2-ketogluconic acid
Galactonate
Galactarate
3-deoxy-Pentitol
Sucrose
Galactitol
Palmitic acid
Mannose
Galacturonate
Xylose
Galactose
Stearic acid
Glycerate
GlcNAc
Lactose
Octadecenoic acid
Fucose
Rhamnose
Butanoic acid
3-Hydroxybutyrate
Succinate
Cholesterol
B Threonine
C Arabinose
D Arabitol
E Gluconate
F Glucuronate
G Galacturonate
H Sorbitol
I Xylose
J Melibiose
K Raffinose
L Glucose
Relative Metabolite Abundance
PBS
MH258
STD
HFD
0.65 X
0.18 X
p=0.219
0.14 X
0.23 X
0.22 X
0.02 X
0.09 X
0.11 X
0 X
0.04 X
0.23 X
0.54 X
0.01 X
0.03 X
0.67 X
p=0.2339
2.16 X
0.81 X

**Figure S7. Dietary context influences the abundance and consumption of key carbon sources during *K. pneumoniae* intestinal colonization. (A)** Absolute signal intensities of selected metabolites measured by GC-MS in cecal contents from antibiotic-treated mice receiving PBS, illustrating the baseline metabolic environment under standard diet (STD) and high-fat diet (HFD) **(B–L)** Relative abundance of the indicated carbon sources in cecal contents from PBS- or MH258-colonized mice maintained on either STD or HFD, as measured by GC- or LC-MS. Comparisons were performed only between PBS- and MH258-colonized mice within the same dietary condition using unpaired two-tailed *t* tests. Fold reductions indicate the decrease in metabolite abundance in MH258-colonized mice relative to the corresponding PBS controls.

Data are presented as means ± SD (*$P < 0.05$; **$P < 0.01$; ***$P < 0.001$; ****$P < 0.0001$).

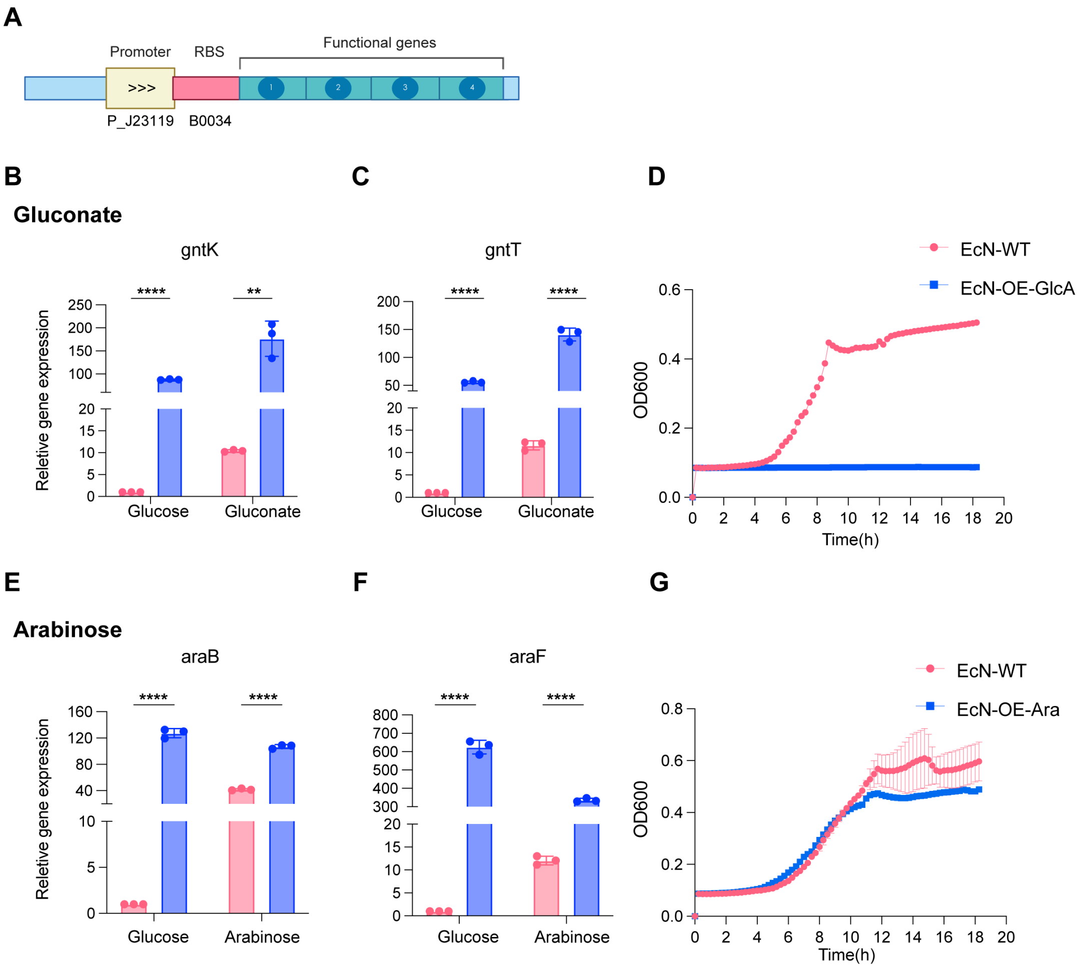


**Figure S8. Overexpression of carbohydrate transporters and metabolic enzymes in *Escherichia coli* Nissle produces unfit strains. (A)** Schematic illustration of the construction of Nissle strains overexpressing carbohydrate transporters and metabolic enzymes. The native promoter was replaced with the strong constitutive promoter J23119, and the native ribosome-binding site (RBS) was replaced with the high-efficiency B0034 RBS. **(B-D)** Overexpression of gluconate utilization genes in *E. coli* Nissle. **(B-C)**, RT–qPCR analysis confirming overexpression of *gntK* and *gntT* in engineered strains grown in M9 minimal medium supplemented with glucose (0.2%) or gluconate (0.2%). Data are shown as means ± SD from three independent experiments, and statistical significance was determined using unpaired two-tailed *t* tests (**P* < 0.05; ***P* < 0.01; ****P* < 0.001; *****P* < 0.0001). **(D)**, growth curves comparing the engineered overexpression strain and the wild-type strain in M9 minimal medium supplemented with gluconate (0.2%). **(E-G)** Overexpression of arabinose utilization genes in *E. coli* Nissle. Experiments were performed as in (B), including RT–qPCR validation of gene (*araB* and *araF*) overexpression and growth curve analysis in arabinose (0.2%) -supplemented M9 medium.

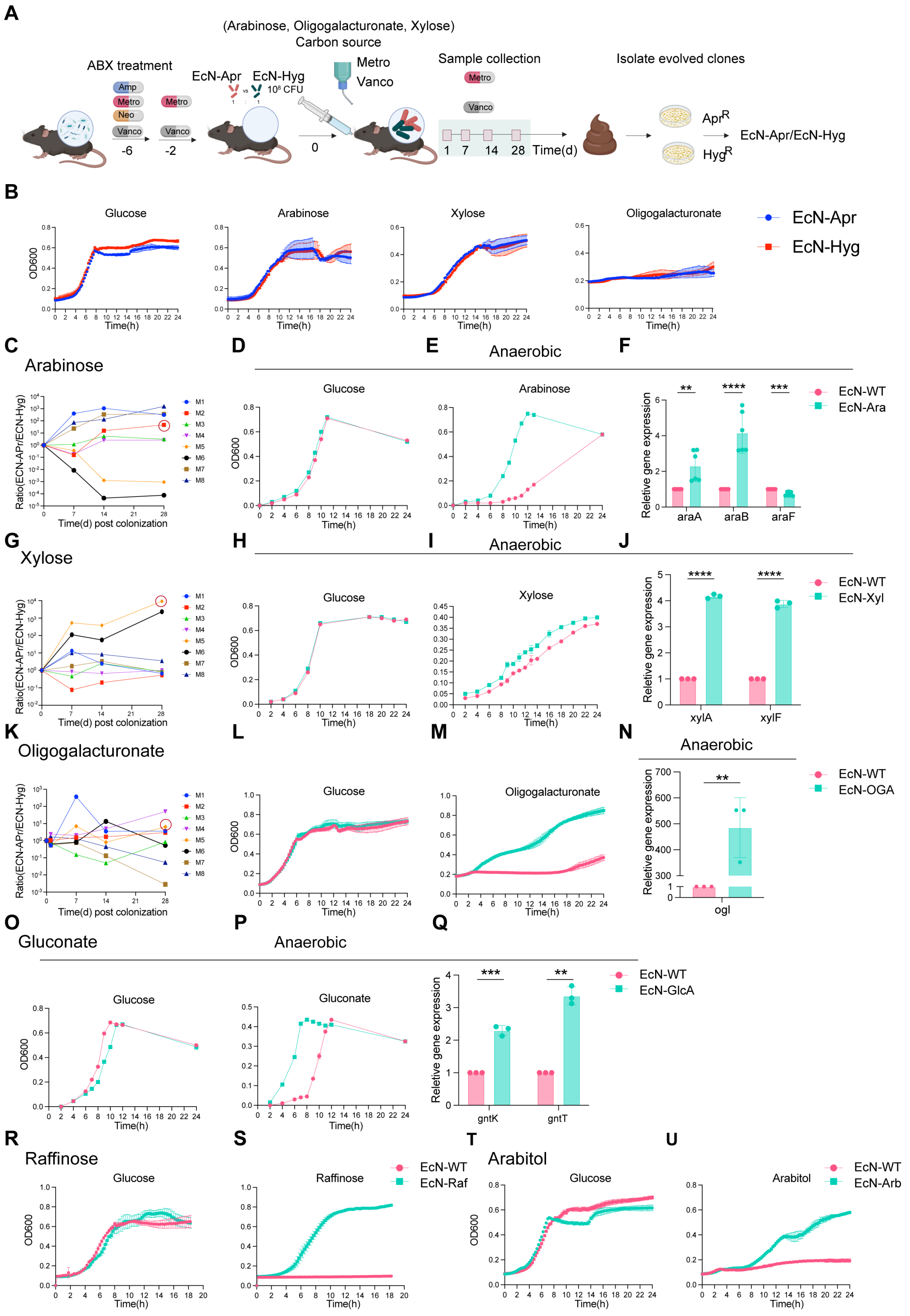


V

| Strain | Mutation type | Effect | Gene | product |
|---|---|---|---|---|
| EcN-OGA | snp | stop_gained c.1571G>A p.Trp524* | malT | Master transcriptional activator of the maltose utilization regulon. |
| | ins | frameshift_variant c.53_54dupAA p.Val19fs | kdgR | Transcriptional repressor of KDG and hexuronate catabolic genes involved in pectin-derived sugar utilization |
| | del | frameshift_variant c.1041delG p.Gly350fs | | MFS transporter |
| EcN-Ara | snp | missense_variant c.843T>A p.Ser281Arg | dacA | D-alanyl-D-alanine carboxypeptidase DacA |
| | snp | missense_variant c.58G>A p.Val20Met | araC | Arabinose operon transcriptional regulator AraC |

**Figure S9. Functional validation of optimized *Escherichia coli* Nissle strains for enhanced utilization of selected carbon sources. (A)** Schematic overview of the *in vivo* experimental evolution strategy used to optimize *Escherichia coli* Nissle (EcN) for improved utilization of key carbon sources during intestinal colonization. For arabinose, xylose, and oligogalacturonate, *in vivo* experimental evolution was performed by introducing two ECN-WT strains carrying neutral chromosomal antibiotic resistance markers (apramycin or hygromycin) at a 1:1 ratio into antibiotic-treated mice supplied with 0.6% of the indicated carbon source in drinking water, followed by longitudinal fecal sampling to identify selectively enriched clones. **(B)** Aerobic growth curves of ECN strains carrying distinct neutral antibiotic resistance markers (EcN-Apr and EcN-Hyg) in M9 minimal medium supplemented with glucose (0.2%), arabinose (0.2%), xylose (0.2%), or oligogalacturonate (0.5%). Growth curve experiments were independently repeated three times, and data are presented as mean ± SEM. **(C-N)** *In vivo* adaptation and functional validation of EcN optimized for arabinose (C-F), xylose (G-J), and oligogalacturonate (K-N) utilization. Temporal dynamics of antibiotic-resistance marker frequency during EcN colonization of the mouse gut when 0.6% arabinose (C), xylose (G), or oligogalacturonate (K) were continuously supplied in the drinking water, revealing reproducible enrichment of adapted EcN variants. Red circles indicate mice/time points from which the evolved clones were selected. Anaerobic or aerobic growth curves comparing optimized EcN isolates recovered from *in vivo* experimental evolution with EcN-WT in M9 minimal medium supplemented with glucose (0.2%) or the corresponding carbon source (arabinose 0.2% (D-E), xylose 0.2% (H-I), or oligogalacturonate 0.5% (L-M)). Growth curve experiments were independently repeated two or three times, and data are presented as mean ± SEM. (F,J,N) RT–qPCR analysis of bacteria harvested at logarithmic phase, showing increased expression of the corresponding carbon metabolism genes in optimized EcN strains relative to EcN-WT. RT–qPCR experiments were performed in three independent biological replicates, with data presented as mean ± SD. Statistical significance was determined using unpaired two-tailed *t* tests (**P* < 0.05; ***P* < 0.01; ****P* < 0.001; *****P* < 0.0001). **(F)** Anaerobic growth and transcriptional profiling of gluconate-optimized EcN strains (Δ*gntR*). Optimized strains and ECN-WT were grown in M9 minimal medium supplemented with glucose (0.2%) or gluconate (0.2%). RNA was extracted at logarithmic phase for RT–qPCR analysis of gluconate-associated metabolic genes. Growth curve experiments were independently repeated two or three times, with data presented as mean ± SEM. RT–qPCR experiments were performed in three independent biological replicates, with data presented as mean ± SD. Statistical significance was determined using unpaired two-tailed *t* tests (**P* < 0.05; ***P* < 0.01; ****P* < 0.001; *****P* < 0.0001). **(R-U)** Aerobic growth curves of raffinose (R-S)- and arabitol (T-U)-optimized EcN strains in M9 minimal medium supplemented with glucose (0.2%) or raffinose (0.2%) and arabitol (0.5%), respectively. Optimization for raffinose was produced by engineering EcN with the MH258 gene *rafB*, *cscA*, and *rafD* and for arabitol with the MH258 gene *dalT*, *dalD*, and *atlB*, in a low-copy, stable plasmid (PACYC177). Growth curve experiments were independently repeated three times, and data are presented as mean ± SEM. **(V)** Whole-genome sequencing analysis of the evolved EcN-OGA and EcN-Ara strains. Mutations identified relative to the parental EcN strain are shown. EcN-OGA contained mutations including disruption of the oligogalacturonate utilization repressor *kdgR*, whereas EcN-Ara harbored a nonsynonymous mutation in the arabinose utilization regulator *araC*, both of which are likely associated with the enhanced utilization of their respective carbon sources.

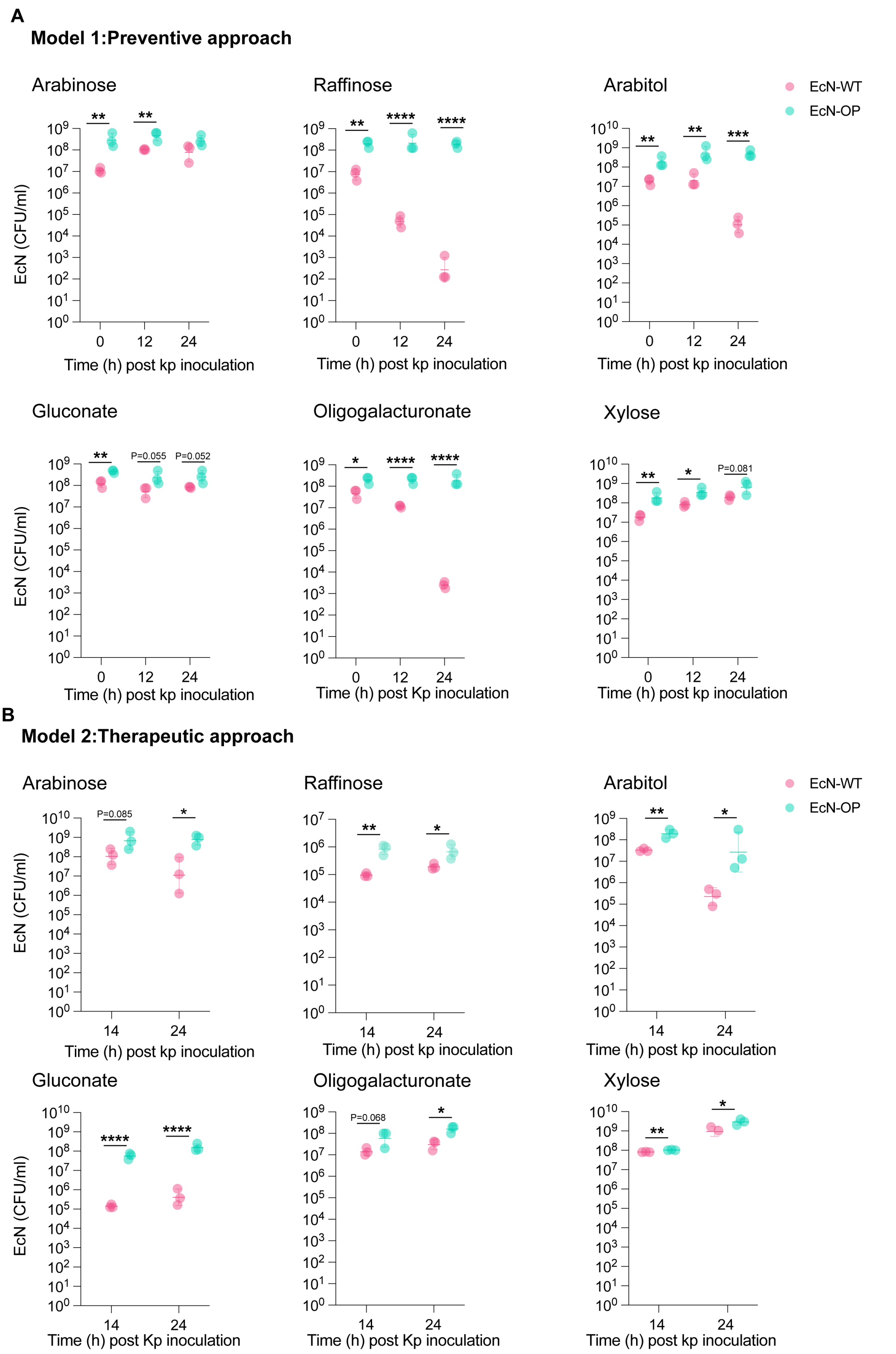


**Figure S10. EcN abundance in *in vitro* protection and decolonization assays of MH258.**

ECN colony-forming units (CFU) quantified in parallel with the *K.pneumoniae* measurements shown in Figure 5E and 5F. **(A)** EcN abundance in the *in vitro* protection model. **(B)** EcN abundance in the *in vitro* decolonization model. Data correspond to the same experimental setup shown in Figure 5F. Each dot represents one biological replicate. Shown is one representative of three independent experiments. Statistical significance was determined using unpaired two-tailed *t* tests (*$P < 0.05$; **$P < 0.01$; ***$P < 0.001$; ****$P < 0.0001$).

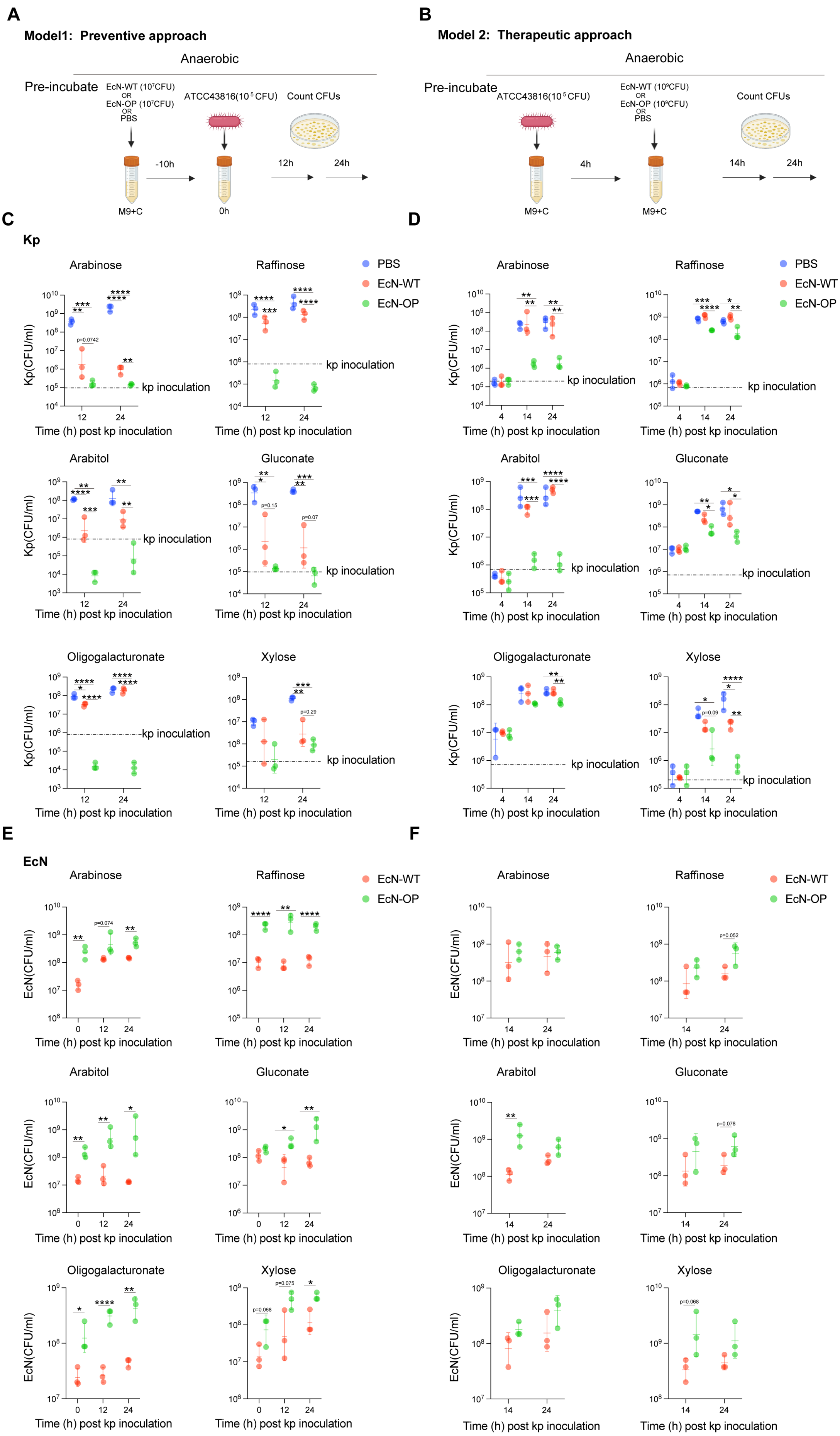
A
Model1: Preventive approach
Anaerobic
Pre-incubate
EcN-WT (10⁷CFU) OR EcN-OP (10⁷CFU) OR PBS
ATCC43816(10⁵CFU)
Count CFUs
-10h
0h
12h
24h
M9+C
B
Model 2: Therapeutic approach
Anaerobic
Pre-incubate
ATCC43816(10⁵CFU)
EcN-WT (10⁸CFU) OR EcN-OP (10⁸CFU) OR PBS
Count CFUs
4h
14h
24h
M9+C
C
Kp
Arabinose
Raffinose
Arabitol
Gluconate
Oligogalacturonate
Xylose
PBS
EcN-WT
EcN-OP
Kp(CFU/ml)
kp inoculation
Time (h) post kp inoculation
D
E
EcN
EcN(CFU/ml)
F

**Figure S11. EcN-OP strains suppress hypervirulent *K. pneumoniae* ATCC43816 expansion *in vitro*. (A)** Schematic of the *in vitro* protection model. Minimal M9 medium supplemented with glucose or the indicated individual carbon source was inoculated with PBS, EcN-WT, or the corresponding EcN-OP strain ($10^7$ CFU). After 10 h of pre-colonization, *K. pneumoniae* ATCC43816 ($10^5$ CFU) was introduced, and bacterial abundance was quantified by selective plating at the indicated time points. **(B)** Schematic of the *in vitro* therapeutic (decolonization) model. *K. pneumoniae* ATCC43816 ($10^5$ CFU) was first inoculated into M9 minimal medium containing glucose or the indicated carbon source. After establishment, EcN-WT, the corresponding EcN-OP strain ($10^9$ CFU), or PBS was added, and bacterial abundance was quantified by selective plating at the indicated time points. **(C, D)** Expansion of *K. pneumoniae* ATCC43816 in the protection (C) and therapeutic (D) models. Dashed lines indicate the initial inoculum level of *K. pneumoniae* ATCC43816. **(E, F)** Growth of EcN-WT and the corresponding EcN-OP strains in the protection (E) and therapeutic (F) models.

Shown is one representative of three independent experiments. Data are presented as geometric means ± geometric SD. Statistical significance for *K. pneumoniae* abundance was determined using one-way ANOVA followed by Tukey's multiple comparisons test. Statistical significance for EcN abundance was determined using unpaired two-tailed *t* tests (**P* < 0.05; ***P* < 0.01; ****P* < 0.001; *****P* < 0.0001).

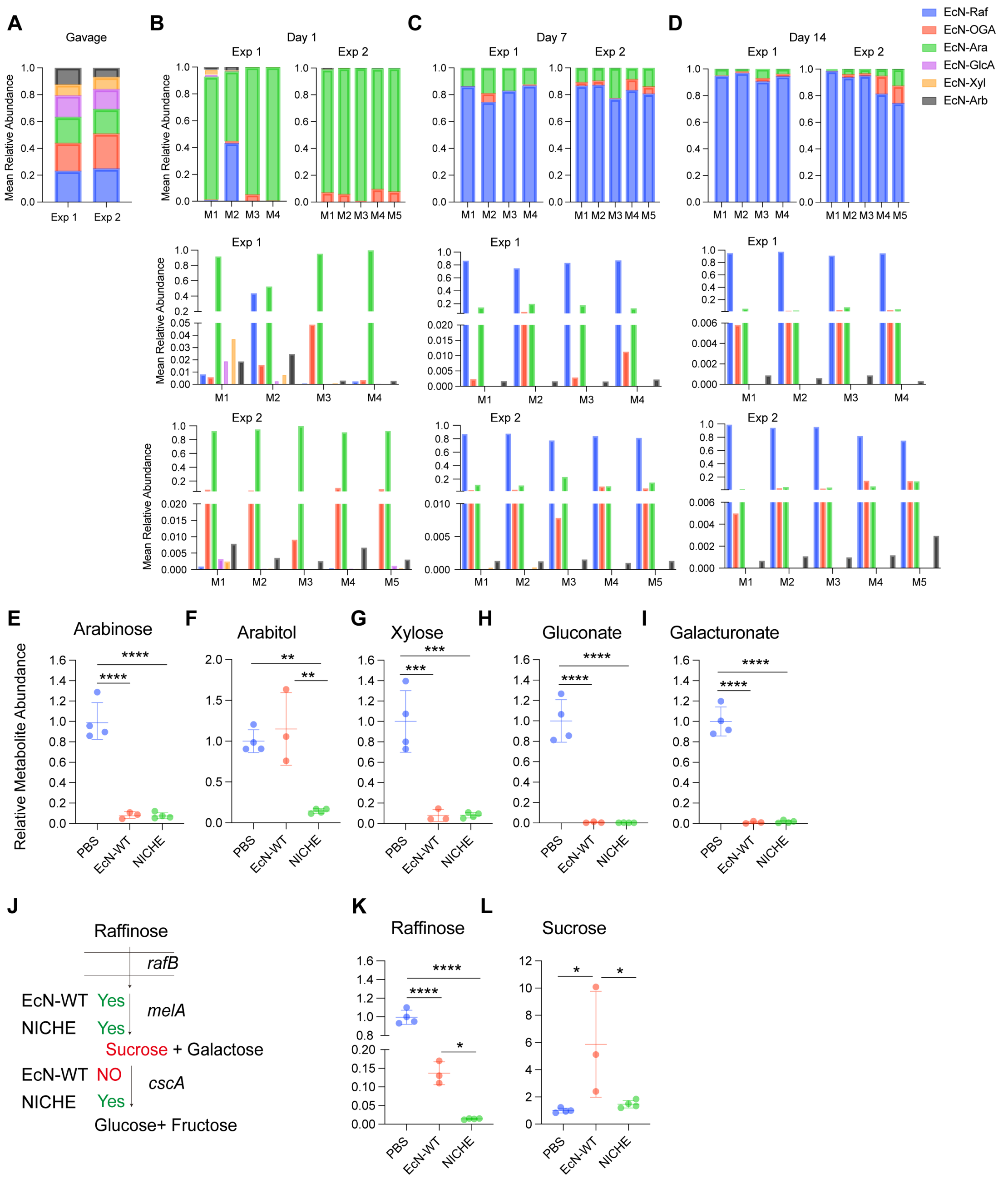


**Figure S12. *In vivo* dynamics of the NICHE consortium and metabolomic characterization of carbon source utilization. (A–D)** Relative abundance of each EcN strain within the NICHE consortium in the inoculum (A) and at days 1 (B), 7 (C), and 14 (D) after intestinal colonization, determined by barcode-specific qPCR. Data are from two independent experiments. **(E–I)** Relative abundances of arabinose (E), arabitol (F), xylose (G), gluconate (H), and glucuronate (I) in cecal contents from mice colonized with EcN-WT or NICHE, determined by GC–MS. **(J)**

Schematic illustrating the proposed raffinose utilization pathway in EcN-WT and NICHE. EcN-WT hydrolyzes raffinose but is unable to further metabolize the resulting sucrose, resulting in sucrose accumulation. In contrast, NICHE is capable of complete raffinose utilization, and consequently no sucrose accumulation was detected. **(K)** Relative abundance of raffinose in cecal contents from mice colonized with EcN-WT or NICHE, determined by LC–MS. **(L)** Relative abundance of sucrose in cecal contents from mice colonized with EcN-WT or NICHE, determined by GC–MS. Data are presented as means ± SD. For metabolomic analyses (E–L), statistical significance was determined using one-way ANOVA followed by Tukey's multiple comparisons test (**P* < 0.05; ***P* < 0.01; ****P* < 0.001; *****P* < 0.0001).

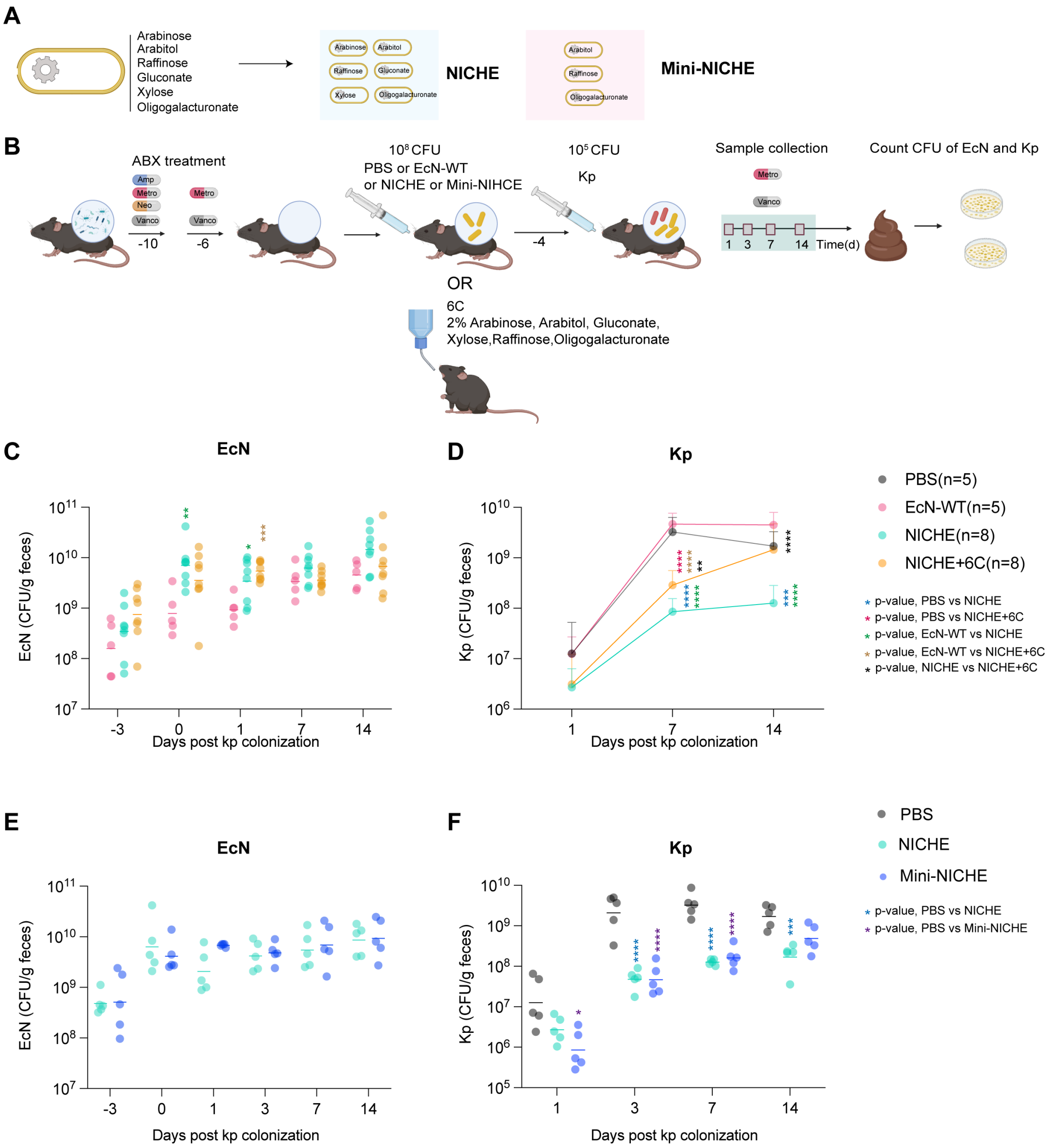


**Figure S13. Evaluation of a minimal NICHE consortium and carbon source supplementation *in vivo*. (A)** Composition of the NICHE and mini-NICHE consortia. NICHE consists of six EcN-OP strains individually optimized for utilization of arabinose, arabitol, raffinose, oligogalacturonate, gluconate, and xylose, whereas mini-NICHE consists of the three EcN-OP strains optimized for utilization of arabitol, raffinose, and oligogalacturonate. **(B)** Schematic of the *in vivo* therapeutic experiment. Antibiotic-treated mice were orally gavaged with NICHE or mini-NICHE. To determine whether restoration of depleted nutrients could reverse NICHE-mediated colonization resistance, an additional NICHE-treated group received drinking water supplemented with a 2% (w/v) mixture of the six target carbon sources (arabinose, arabitol, raffinose, oligogalacturonate, gluconate, and xylose). Four days later, mice were challenged with multidrug-resistant *K. pneumoniae* MH258, and fecal samples were collected at the indicated time points for CFU enumeration. **(C,D)** Fecal abundance of EcN (E) and *K. pneumoniae* MH258 (F) following therapeutic administration of PBS, EcN-WT, NICHE, or NICHE supplemented with a 2% (w/v) mixture of the six target carbon sources in the drinking water. Carbon source supplementation restored *K. pneumoniae* colonization, consistent with reversal of the colonization resistance mediated by NICHE through depletion of these nutrients. **(E,F)** Fecal abundance of EcN (C) and *K. pneumoniae* MH258 (D) following therapeutic administration of PBS, NICHE, or mini-NICHE. Mini-NICHE, consisting of the three dominant EcN-OP strains (EcN-Arb, EcN-Raf, and EcN-OGA), conferred colonization resistance comparable to that of the full NICHE consortium.Data shown are pooled from two independent experiments, and are presented as geometric means ± geometric SD. Statistical significance was determined using one-way ANOVA followed by Tukey's multiple comparisons test (*$P < 0.05$; **$P < 0.01$; ***$P < 0.001$; ****$P < 0.0001$).

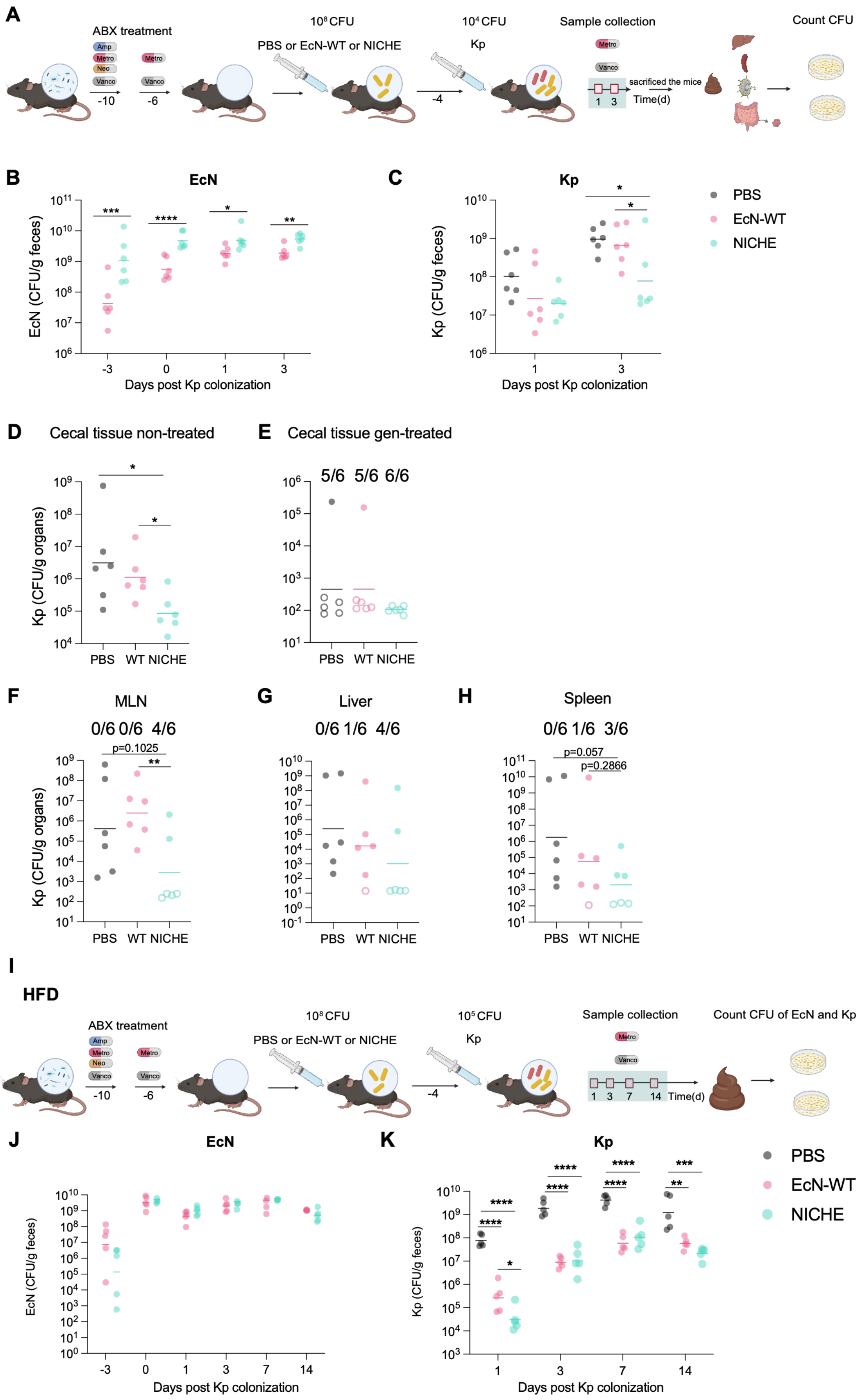

A
ABX treatment
Amp
Metro
Neo
Vanco
-10
-6
$10^8$ CFU
PBS or EcN-WT or NICHE
$10^4$ CFU
Kp
-4
Sample collection
sacrificed the mice
1 3
Time(d)
Count CFU
B
EcN
EcN (CFU/g feces)
Days post Kp colonization
C
Kp
Kp (CFU/g feces)
Days post Kp colonization
PBS
EcN-WT
NICHE
D
Cecal tissue non-treated
Kp (CFU/g organs)
PBS WT NICHE
E
Cecal tissue gen-treated
5/6 5/6 6/6
PBS WT NICHE
F
MLN
0/6 0/6 4/6
p=0.1025
Kp (CFU/g organs)
PBS WT NICHE
G
Liver
0/6 1/6 4/6
PBS WT NICHE
H
Spleen
0/6 1/6 3/6
p=0.057
p=0.2866
PBS WT NICHE
I
HFD
ABX treatment
$10^8$ CFU
PBS or EcN-WT or NICHE
$10^5$ CFU
Kp
Sample collection
1 3 7 14 Time(d)
Count CFU of EcN and Kp
J
EcN
EcN (CFU/g feces)
Days post Kp colonization
K
Kp
Kp (CFU/g feces)
Days post Kp colonization
PBS
EcN-WT
NICHE

(A) Schematic of the in vivo protection model using hypervirulent *K. pneumoniae* ATCC43816. Antibiotic-treated mice were orally gavaged with PBS, EcN-WT, or NICHE. Four days later, mice were challenged with ATCC43816 (10^4 CFU). Fecal samples were collected at the indicated time points. On day 3 after challenge, mice were euthanized, and cecal tissue, mesenteric lymph nodes (MLNs), liver, and spleen were collected for CFU enumeration. (B, C) Fecal burdens of ATCC43816 (B) and EcN (C) following challenge. (D, E) ATCC43816 burdens in total cecal tissue (D) and gentamicin-protected cecal tissue (E). (F–H) ATCC43816 burdens in mesenteric lymph nodes (F), liver (G), and spleen (H). Open symbols indicate samples in which no colonies were detected, whereas filled symbols indicate samples with detectable CFU. Because bacterial burdens are expressed as CFU per gram of tissue, the limit of detection (LOD) varies among individual samples depending on the weight of tissue processed. Numbers above the graphs indicate the number of mice with bacterial burdens below the LOD. Data shown are from one representative experiment (n = 6 mice per group). Data are presented as geometric means ± geometric SD. **(I)** Schematic of the in vivo protection experiment performed under HFD conditions using multidrug-resistant *K. pneumoniae* MH258. Antibiotic-treated HFD-fed mice were orally gavaged with EcN-WT or NICHE. Four days later, mice were challenged with MH258 ($10^5$ CFU), and fecal samples were collected longitudinally for CFU enumeration. **(J, K)** Fecal burdens of EcN (J) and MH258 (K) in HFD-fed mice treated with EcN-WT or NICHE. Data shown are from one representative experiment (n = 5 mice per group). Data are presented as geometric means ± geometric SD. For *K. pneumoniae* abundance, comparisons among treatment groups at each time point were performed using one-way ANOVA followed by Tukey's multiple comparisons test. For EcN abundance, comparisons between two groups were performed using unpaired two-tailed *t* tests (*$P < 0.05$; **$P < 0.01$; ***$P < 0.001$; ****$P < 0.0001$).
